# RADAR: Differential analysis of MeRIP-seq data with a random effect model

**DOI:** 10.1101/867903

**Authors:** Zijie Zhang, Qi Zhan, Mark Eckert, Allen Zhu, Agnieszka Chryplewicz, Dario F De Jesus, Decheng Ren, Rohit N Kulkarni, Ernst Lengyel, Chuan He, Mengjie Chen

**Affiliations:** Section of Genetic Medicine, Department of Medicine, Department of Human Genetics, The University of Chicago, Chicago, IL 60637, USA; Department of Chemistry, Department of Biochemistry and Molecular Biology, and Institute for Biophysical Dynamics, The University of Chicago, Chicago, IL 60637, USA; Howard Hughes Medical Institute, The University of Chicago, Chicago, IL 60637, USA; Department of Obstetrics & Gynecology, Section of Gynecologic Oncology, The University of Chicago, Chicago, IL 60637, USA; Section of Islet Cell and Regenerative Biology, Joslin Diabetes Center and Harvard Medical School, Boston, MA 02215 USA; Section of Endocrinology, Diabetes & Metabolism, Department of Medicine, The University of Chicago, Chicago, IL 60637 USA

**Keywords:** *N*^6^-adenosine methylation (m^6^A), differential methylation, MeRIP-seq

## Abstract

Epitranscriptome profiling using MeRIP-seq is a powerful technique for in vivo functional studies of reversible RNA modifications. We develop RADAR, a comprehensive analytical tool for detecting differentially methylated loci in MeRIP-seq data. RADAR enables accurate identification of altered methylation sites by accommodating variability of pre-immunoprecipitation expression level and post-immunoprecipitation count using different strategies. In addition, it is compatible with complex study design when covariates need to be incorporated in the analysis. Through simulation and real datasets analyses, we show that RADAR leads to more accurate and reproducible differential methylation analysis results than alternatives, which is available at https://github.com/scottzijiezhang/RADAR.

## Background

The RNA epigenetics gold rush in the past few years has brought reversible RNA modifications into the spotlight as an important mechanism of gene regulation. Particularly, *N*^6^-adenosine methylation (m^6^A), the most abundant modification on mRNA, has drawn extensive attention due to its important functions in various biological systems [1–3]. Methylated RNA immunoprecipitation sequencing (MeRIP-seq) [4–6] is a key technique used in mRNA modification studies that has enabled us to survey the epitranscriptome using various study designs: 1) identifying the location of modification on the transcript by performing peak calling on MeRIP-seq samples of certain phenotype or experimental condition and, 2) identifying differentially methylated loci by comparing MeRIP-seq samples across different phenotypical or experimental groups. These analyses connect mRNA modifications with phenotypes and have the great potential to reveal its functional consequences.

Early studies performing qualitative analysis compared peaks called in one experimental group versus peaks in another group and identified peaks unique to each experimental group as differentially methylated peaks. However, many differential peaks identified by this method are caused by boundary cases at the peak-detection threshold rather than true present/absent of peaks as noted in an recent study [7]. To enable cross-group comparisons, a few methods have been developed and applied to analyze differential methylation in MeRIP-seq data [8–11]. ExomePeak uses Fisher’s exact test for differential methylation identifications and its later version uses a likelihood ratio test based on the binomial distribution (termed “bltest”) [11, 12]. MeTPeak uses a beta-binomial model to infer differential peaks [8]. DRME and its improved version — QNB uses a model based on the negative binomial distribution [9, 10].

While existing methods have yielded promising results, they also have important drawbacks: (1) Current methods [8–11] designed for small sample size scenario ignore the existence of confounding factors and cannot accommodate complex study designs with covariates (such as age, gender, etc.) that are frequently encountered in patient or animal studies with larger sample sizes. (2) Most of the differential gene expression (DE) analysis tools such as edgeR, DESeq2 and Sleuth [13–15] are compatible with complex study designs. But they rely on models developed for RNA-seq experiment and cannot accommodate unique features of MeRIP-seq data. A standard MeRIP-seq experiment yields an INPUT and an Immunoprecipitation (IP) library for each sample. The INPUT library is the initial RNA fragments pool prior to the antibody pull-down — a measurement of RNA expression level. The IP library represents the RNA fragments carrying modified bases captured by antibody pull-down — a measurement of methylated RNA abundance. RNA Differential Methylation (DM) is defined as the alteration of methylated RNA abundance conditioning on the RNA expression background. Thus, DM analysis requires assessment of RNA methylation change based on pre-IP and post-IP measurements in pairs. In contrast, DE analysis tools only compare a single read count measurement across samples. (3) Current MeRIP-seq specific tools [8, 9, 11] use peak (~250bp) read counts in the INPUT library as measurement of RNA expression for a gene (~11kb). However, the variability in a small genomic range across samples due to the sparsity of reads sampled can result in unwanted variation to the expression level estimation if using local read counts. QNB combines local read counts of both INPUT and IP as an estimator of the expression level to mitigate this problem. However, incorporating IP read counts can cofound pre-IP expression level with post-IP RNA abundance, leading to biased estimation of expression level. Inaccurate expression level measurement can lead to substantial false discoveries in the subsequent DM analysis. Thus, the utilization of INPUT library to account for pre-IP RNA expression level needs to be further optimized.

To combat these challenges and allow for accurate identification of differentially methylated loci, we present a novel approach to perform **R**NA methyl**A**tion **D**ifferential **A**nalysis in **R** (**RADAR**) for MeRIP-seq data. RADAR accounts for variation in pre-IP RNA and in post-IP read counts using different strategies. Specifically, RADAR uses gene-level read counts instead of peak-level read counts in the INPUT library as a robust measurement of the initial pre-IP RNA expression level. In addition, RADAR uses a flexible Poisson random effect model to accommodate over-dispersion in the post-IP read counts due to variability of biological replicates and noise introduced in the immunoprecipitation process. This generalized linear model framework enables incorporation of covariates in complex study designs.

We benchmarked the performance of RADAR with alternative methods on simulated data by different data generating models. We showed RADAR achieved higher sensitivity and specificity compare to existing alternative methods. We also demonstrated the performance of RADAR on real MeRIP-seq data by applying it to four high quality m^6^A-meRIP-seq (aka m^6^A-seq) datasets generated by us and others, including an ovarian cancer dataset (GSE 119168) consisting of 7 normal fallopian tubes tissue from healthy individuals and 6 metastatic omental tumors, a Type 2 Diabetes (T2D, GSE 120024) dataset consisting of human islets from 8 type II diabetes patients and 7 healthy control patients with samples being processed in three batches due to different sample acquisition times, a mice liver (GSE 119490) dataset consisting of mouse liver from 4 wild type mice and 4 METTL14 knockout mice and a mice brain (GSE 113781) dataset consisting of 7 mouse cortex samples of stress exposed mice and 7 from control mice. We showed that our approach can accommodate distinct study designs and led to more sensitive and reproducible DM loci identification than possible alternatives.

## Results

### RADAR overcomes challenges in modeling MeRIP-seq data and accommodates complex study designs

Using BAM files as the input, RADAR first divides transcripts (concatenated exons) into 50-bp consecutive bins and quantifies pre-IP and post-IP read counts for each bin (**Fig. 1A**). Unlike current differential methylation analysis methods [8–11] that scale to library sizes as a way of normalization, which can be strongly skewed by highly expressed genes [16] (**Additional file 1: Fig. S1**), RADAR uses the median-of-ratio method [17] implemented in DEseq2 to normalize the INPUT library for the sake of robustness. For the IP library, RADAR normalizes the fold enrichment computed from the IP counts divided by the INPUT counts, which takes both IP efficiency and IP library size variation into account.

**Figure 1.**
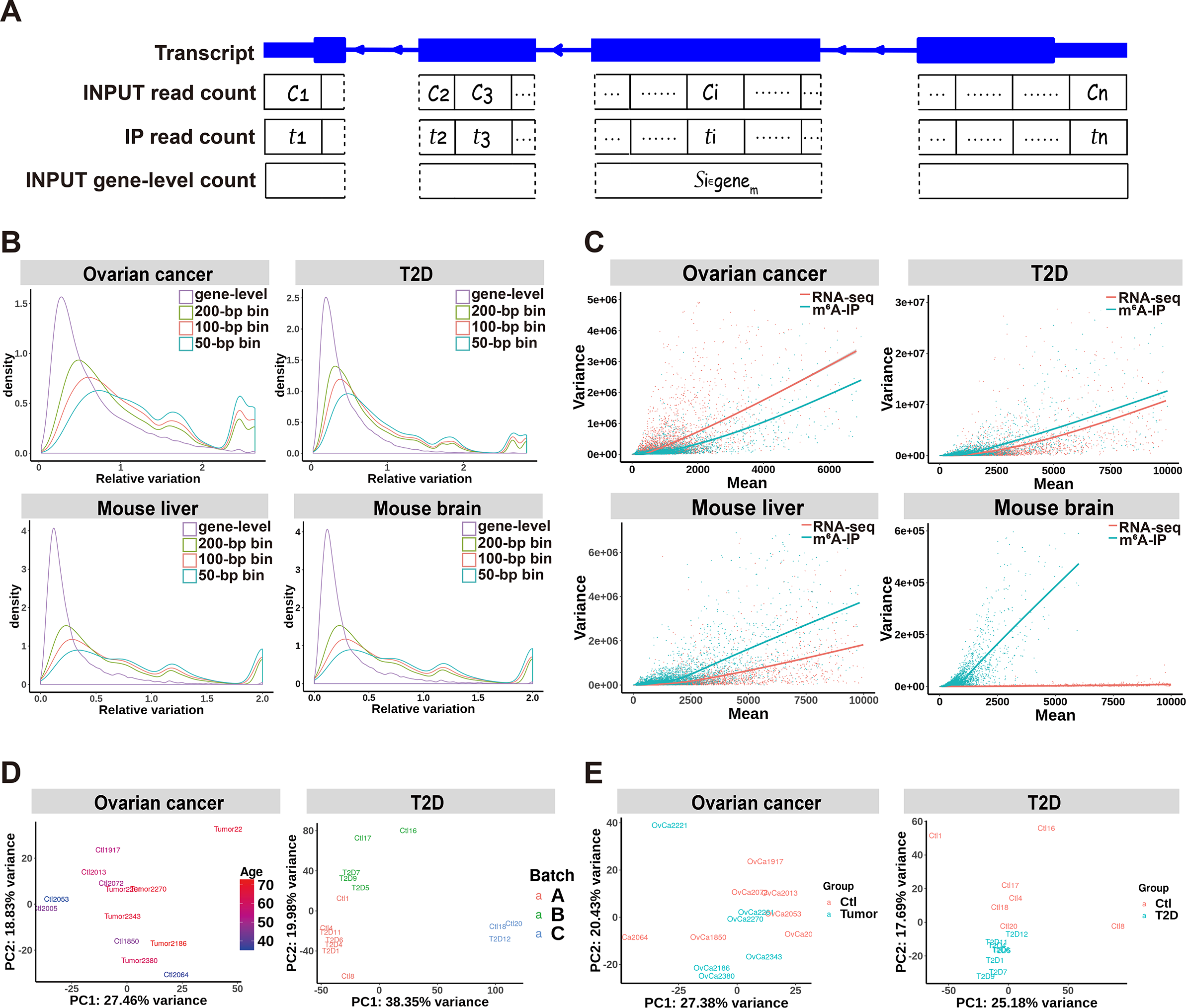
Unique features of m^6^A-seq (MeRIP-seq) data. RADAR divides concatenated exons of a gene into consecutive bins and models the immunoprecipitation (IP)-enriched read counts in such bins. (**a**) depicts a pair of read counts in the INPUT and the IP library in the *i*-th bin as *C*_*i*_ and *t*_*i*_. In the RADAR workflow, the gene-level read count of the input library 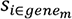 substitutes the bin-level read count *C*_*i*_ as the representation of the pre-IP RNA levels of the *i*-th bin. (**b**) compares the relative variation of gene-level and bin-level (local) read counts of different bin sizes in four m^6^A-seq datasets, suggesting that unwanted variation can be reduced using gene-level counts as the estimates of pre-IP RNA levels. Panel (**c**) compares the cross-sample mean and variance of regular RNA-seq (pre-IP counts) and m^6^A-seq (post-IP read counts adjusted for pre-IP RNA level variation) data in four m^6^A-seq datasets. The fitted curvature of m^6^A-seq can differ from that of RNA-seq, indicating that m^6^A-seq may have a different mean-variance relationship from RNA-seq. Biological and experimental confounding factors are often encountered in patient samples. (**d**) shows the first two principal components (PCs) of m^6^A enrichment in each dataset, where the samples are colored by covariates that need to be accounted for. m^6^A enrichment was represented by IP sample read counts adjusted for pre-IP (INPUT) RNA level variation. (**e**) shows the first two PCs after regressing out known covariates — age in the ovarian cancer dataset and batch in the T2D dataset. After regressing out the covariate, samples are separated by disease conditions on the PCA plot.

After proper normalization across all samples, RADAR then calculates the methylation level for each bin conditioned on its pre-IP RNA expression level for each sample. In contrast to previous methods [8–11] that use peak-level read counts in the INPUT library as its measurement of pre-IP RNA expression level, we use gene-level read counts as a more robust representation, which is defined as the total number of reads across all bins that span the same gene **(Fig. 1A)**. This choice is motivated by the observation that the median read coverage within each peak is very low —18 reads per peak (7 reads in a 50-bp bin) (**Additional file 1: Fig. S2**) in a typical MeRIP-seq input sample of 20 million (mappable) reads (**Additional file 1: Fig. S3**). Over dispersion of low counts due to random sampling in the sequencing process can introduce substantial unwanted variation to the estimation of pre-IP RNA level. This can be further exacerbated by the uneven distribution of reads caused by local sequence characteristics such as GC content and mappability. Using gene-level counts as the estimate of pre-IP RNA expression level can mitigate the dispersion by increasing the number of reads (272 reads on average) and simultaneously diminishing the effects of sequence characteristics within a gene (**Fig. 1A**). By comparing the variance of read counts across replicates at the gene-level with that at the bin-level, we show that the cross-sample variance is much less at the gene-level than at the bin-level in all three datasets. (**Fig. 1B**).

RADAR models the read count distribution using a Poisson random effect model instead of a negative binomial distribution, which is commonly used in RNA-seq analysis [13, 15, 17] as well as in DRME and QNB for MeRIP-seq analysis [9, 10]. Negative binomial distribution-based models assume a quadratic relationship between mean read counts and their variance across all genes. We observe in real m^6^A-seq datasets that the mean-variance relationship of post-IP counts across genes significantly differs from that of regular RNA-seq counts (i.e., pre-IP counts). The former does not always follow a similar quadratic curvature and can exhibit very different patterns of variability (**Fig. 1C**, **Additional file 1: Fig. S4**). To overcome these limitations, RADAR applies a more flexible generalized linear model framework (**see Method**) that captures variability through random effects.

Another important advancement of RADAR, compared to existing MeRIP-seq data analysis tools [8–11], is the flexibility to incorporate covariates and permit more complex study design. Phenotypic covariates such as age and gender as well as experimental covariates such as batch information are often encountered in epitranscriptomic profiling studies with heterogenous patient samples. Covariates such as litter and age are common in experimental animal studies. For example, in the ovarian cancer dataset, the age of the tissue donors is partially confounded with predictor variable – disease status. In the T2D islets dataset, the variance of the first two principal components is confounded with the sequencing batch (**Fig. 1D**). After regressing out the batch effect, the remaining variance can be better explained by disease status (**Fig. 1E**). This indicates the importance of controlling for potential confounding factors when performing differential methylation tests. The generalized linear model framework in RADAR allows the inclusion of covariates and offers support for complex study designs.

### Comparative benchmarks of different methods using simulated datasets

To evaluate the performance of RADAR in comparison to current methods, we applied RADAR and other methods for MeRIP-seq differential analysis including exomePeak, Fisher’s exact test, MeTDiff and QNB on simulated datasets. We considered four scenarios: the proposed random effect model with/without covariates and the quad-negative binomial (QNB) model adopted from QNB [9] [10] with/without covariates. For each scenario, we evaluated the sensitivity and false discovery rate (FDR) of different methods using ten simulated copies. We first simulated a dataset of 8 samples using the random effect model (Method Section Equation (1), denoted as the simple case). The INPUT library was directly drawn from the T2D dataset. We simulated IP read count adjusted for pre-IP expression level of each bin according to Equation (1) where μ is equal to mean log read count in the “control” group of T2D dataset. The final IP read counts were obtained by rescaling simulated data by the average IP/INPUT ratio observed in the T2D data. In total, we simulated three datasets of 26324 sites in which 20% of sites are true positives with effect size of 0.5, 0.75 or 1, respectively.

For DM loci with an effect size of 0.5, RADAR achieved 29.1% sensitivity and 12.0% FDR at an FDR cutoff of 10%. At the same cutoff, exomePeak and Fisher’s test achieved 72.8% sensitivity/52.5% FDR and 72.2% sensitivity/50.5% FDR, respectively. MeTDiff achieved 10.5% sensitivity and 16.2% FDR. QNB, on the contrary, did not own any power for the small effect size. When the effect size increased, RADAR achieved much higher sensitivity, 77.8% for an effect size of 0.75 and 95.7% for an effect size of 1, while FDR were well calibrated at 10.4% and 10.1%, respectively. exomePeak and Fisher’s test both achieved 89% and 96% sensitivity for effect sizes of 0.75 and 1, respectively, but at the cost of unsatisfactory FDRs, which were greater than 46%. MeTPeak exhibited well-calibrated FDR (12.3% and 11.4%) and moderate sensitivity of 50.4% and 81.5% for effect sizes of 0.75 and 1, respectively. QNB only had low power for an effect size of 1 (beta=1, 13.9% sensitivity and 0.5% FDR). Overall, for the simple case without covariates, RADAR achieved high sensitivity while maintained low FDR at varying true effect sizes (**Fig. 2A**). We then applied the above analysis at varying FDR cutoff and found RADAR achieved the highest sensitivity at a fixed level of empirical FDR (**Additional file 1: Fig S5A**). We note exomePeak and Fisher’s test achieved high sensitivity at all effect sizes as combining read counts across replicates of the same group helped to gain power. As a tradeoff, failing to account for within-group variability resulted in high FDR. On the contrary, RADAR and MeTDiff exhibited well-calibrated FDR while achieved high sensitivity at same levels as exomePeak for large effect sizes. QNB was overconservative and possessed little power.

**Figure 2.**
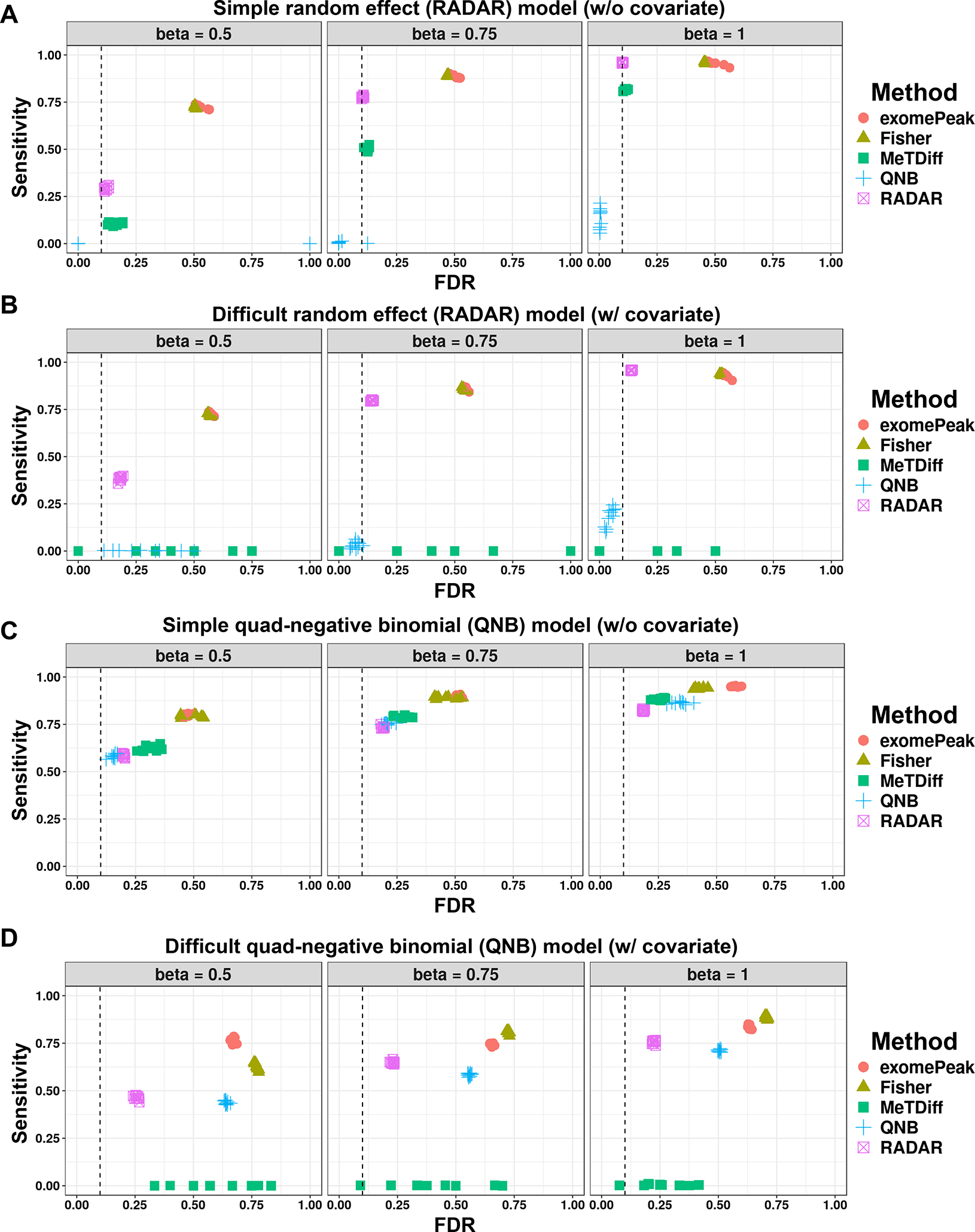
Benchmarking RADAR on two simulation models. We benchmarked RADAR and other alternative methods on simulated data. Using two simulation models — a random effect (RADAR) model and a quad-negative-binomial (QNB) model, we simulated dataset of 8 replicates of varying true effect size (0.5, 0.75 and 1) with and without covariates. We tested different methods on simulated dataset and compared the results at an FDR cutoff of 0.1 with simulated true sites. We show the sensitivity (fraction of true sites detected by the method at an FDR cutoff of 0.1) and false discovery rate (fraction of detected differential sites that are not true sites) of each method applied on data simulated by the random effect model without covariates (**a**) and with covariates (**b**), the quad-negative-binomial model without covariates (**c)** with covariates (**d**), respectively. The FDR cutoff used to select DM sites is labeled by a dashed line.

We next applied the aforementioned methods to the proposed model with a covariate (effect size equal to 2, denoted as the difficult case) (**Fig. 2B**). As a result, at an FDR cutoff of 10%, RADAR achieved 38.4%, 79.7% and 95.7% sensitivity with empirical FDRs slightly higher than those in the simple case (18.2%, 14.4% and 13.7% for effect sizes of 0.5, 0.75 and 1, respectively). MeTDiff, with similar performance as RADAR in the simple case, lost power in the difficult case due to incapability of accounting for confounding factors. exomePeak, Fisher’s test and QNB behaved similarly as in the simple case. The advantage of RADAR over other methods is robust to the choice of FDR cutoff as shown in **Additional file 1: Fig. S5B**. In summary, RADAR outperformed existing alternatives in both cases.

Taking the covariate model with a DM effect size of 0.75 as an example, we also checked the distributions of effect size estimates and p-values obtained from each method. In all methods, effect sizes were overall correctly estimated with estimates for “true” sites centered at 0.75 (**Additional file 1: Fig. S6A**) and that for null sites centered at zero (**Additional file 1: Fig. S6B**). However, we note the distribution of beta estimates is narrower for RADAR, especially in the difficult case, suggesting a more confident estimation. P-values of exomePeak and Fisher’s test at null sites were enriched near zero, indicating over-detection of false positive signals (**Additional file 1: Fig. S6C**). We also observed many large p-values obtained by QNB for “true” sites in both cases and MeTDiff in the difficult case, which suggested a high false negative rate (**Additional file 1: Fig. S6D**).

We then repeated simulation studies using the QNB model. Instead of setting the variances of INPUT and IP libraries equal as presented in the QNB paper, we let the variance of IP read count be larger than that of INPUT. This setting better reflects our observation in the real data as extra noise can be introduced during immunoprecipitation process for IP reads generation (**Additional file 1: Fig. S4**). In the simple case without covariates, RADAR exhibited the lowest empirical FDR (18.9% and 18.5%) despite of slightly lower sensitivity comparing to other methods (73.5% and 82.3%) when the effect sizes were relatively large (for effect sizes of 0.75 and 1). QNB performed better when the effect size was small with 58.6% sensitivity and 15.6% FDR for an effect size of 0.5 (**Fig. 2C**). The results were consistent when we evaluated their performance with different FDR cutoffs. Overall, QNB performed slightly better than RADAR with an effect size of 0.5. RADAR achieved similar sensitivity but better calibrated FDR when effect sizes equal to 0.75 and 1 (**Additional file 1: Fig. S5C**). In the model with covariates, RADAR exhibited the lowest empirical FDR, with 25.8%, 23.0% and 22.5% at effect sizes of 0.5, 0.75 and 1, respectively, while other methods either failed to detect the signal or had a higher empirical FDR. Specifically, MeTDiff had sensitivity below 0.5% at varying effect sizes and QNB reached FDRs of 64.1%, 55.8% and 50.5% for effect sizes of 0.5, 0.75 and 1, respectively, at an FDR cutoff of 10% (**Fig. 2D**). The advantage of RADAR over alternative methods hold in the difficult case at varying cutoffs (**Additional file 1: Fig. S5D**). In summary, RADAR outperformed other existing methods in most scenarios, particularly when covariates were present.

### Comparative benchmarks of different methods using four real m^6^A-seq datasets

Next, we compared the performance of different methods using four real m^6^A-seq datasets: ovarian cancer (GSE119168), T2D (GSE120024), mouse liver (GSE119490) and mouse brain (GSE113781). To evaluate the sensitivity of different methods, we first checked the distributions of p-values obtained from corresponding DM tests (**Fig. 3**). In the ovarian cancer, T2D and mouse liver data, Fisher’s test and exomePeak detected the most signals as the p-values are most dense near zero. In these three datasets, RADAR also returned a desirable shape for the p-values histogram in which p-values were enriched near zero while uniformly distributed elsewhere. MeTDiff returned a desired shape only in the ovarian cancer and mouse liver datasets. QNB were overconservative in the ovarian cancer and T2D dataset. All methods failed to return enriched p-values near zero for the mouse brain dataset, suggesting there was no or little signal in this dataset. This is consistent with the original publication that very few differential peaks were detected in this study [7].

**Figure 3.**
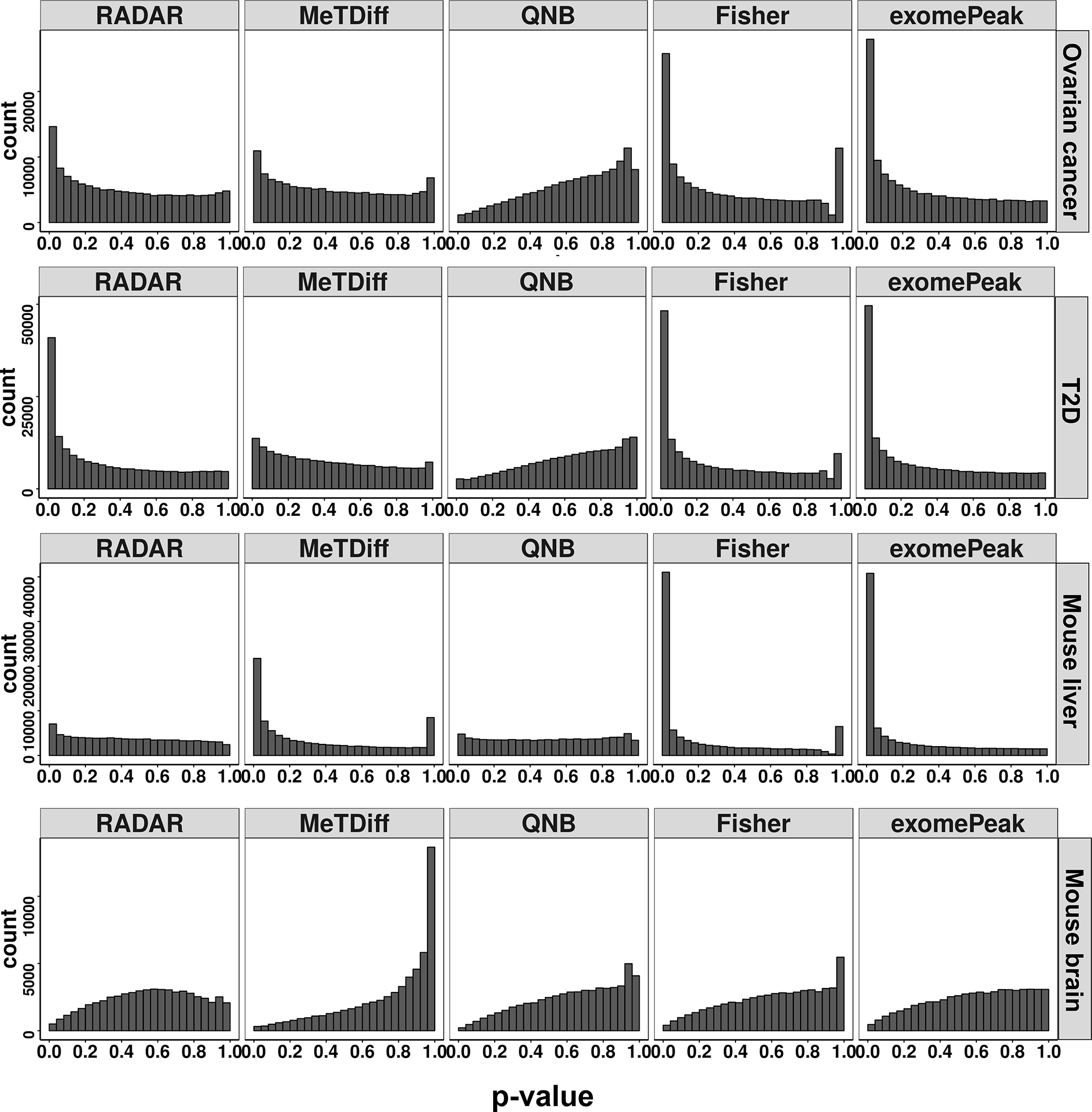
Sensitivity of benchmarked methods on real m^6^A-seq data. We benchmarked RADAR and other alternative methods on four m^6^A-seq data with different characteristics. Each panel shows the histogram of p-values obtained from DM tests using RADAR, MeTDiff, QNB, Fisher’s exact test and exomePeak on each dataset, respectively.

To ensure that well-performed methods achieved high sensitivity while maintain a low FDR, we further performed permutation analyses to obtain the null distribution of p-values for each dataset. Specifically, we shuffled the phenotype labels of samples such that the new labels were not associated with the true ones or any other important confounding factors. We expected the p-values from a permutation test to follow a uniform distribution and the enriched p-values near zero would be considered as false discoveries. For each dataset, we combined test statistics from 15 permuted copies and compared their distribution with the original tests (**Fig. 4**). P-values from Fisher’s test and exomePeak were strongly enriched near zero and only slightly lower than those from the original tests. This suggests the strong signals detected by these two methods are likely to be false discoveries, consistent with the conclusion from simulation analysis. On the contrary, the histograms of p-values from RADAR were close to flat in all datasets, indicating that strong signals detected by RADAR were more likely to be true. MeTDiff exhibited well-calibrated p-values in the ovarian cancer and T2D data but enriched for small p-values in the mouse liver data with an indicated high FDR. QNB test returned conservative p-value estimates in all datasets. Taking together these analyses, we demonstrated that RADAR outperforms the alternatives by achieving high sensitivity and specificity simultaneously in real datasets.

**Figure 4.**
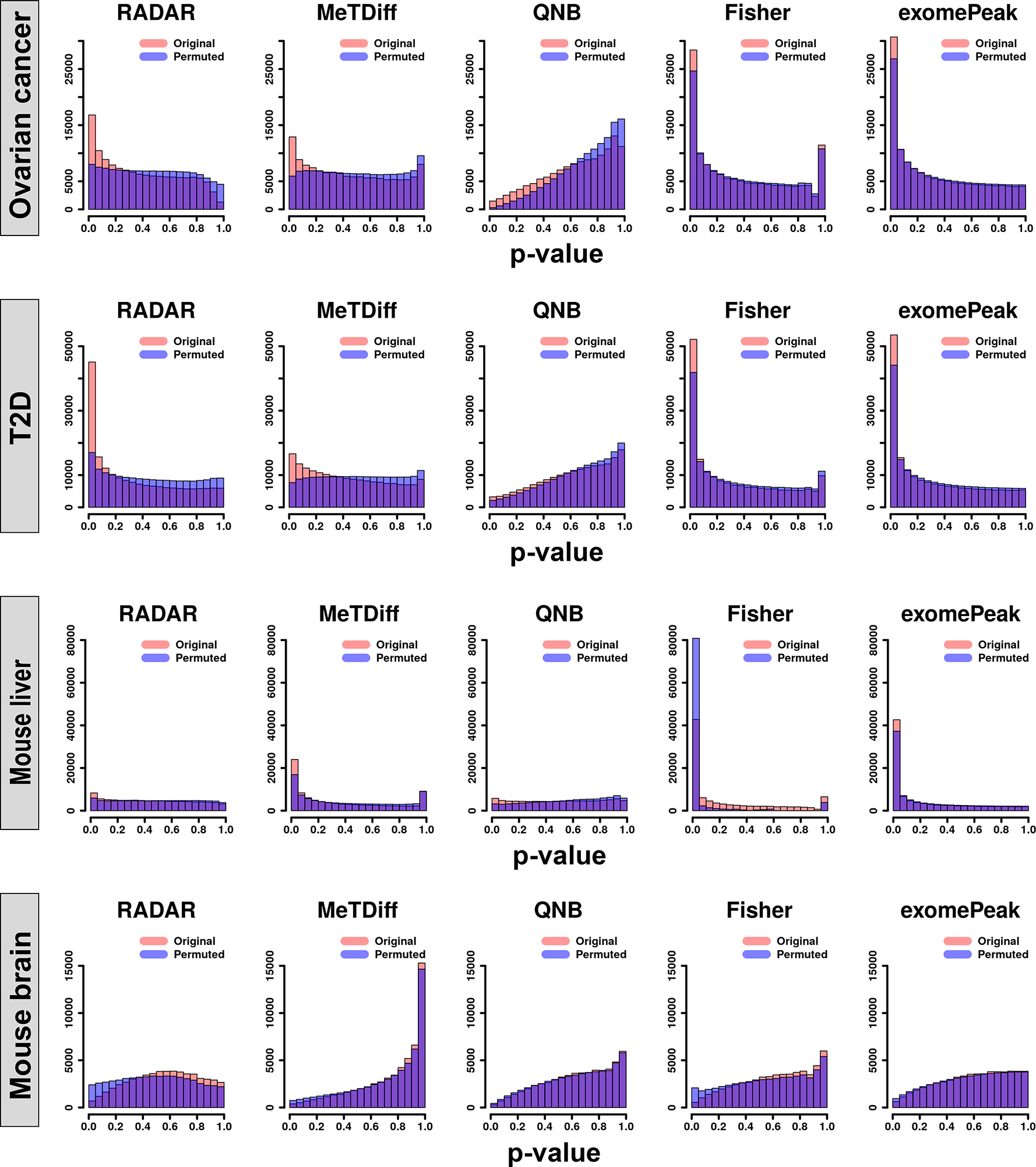
Benchmarking false positive signals using permutation analysis on real m^6^A-seq data. To assess empirical FDR of the test, we permuted the phenotype labels of samples so that the new labels were not associated with true ones. Each panel shows the histograms of p-values obtained from DM tests on 15 permuted copies (blue) and those from the tests on the original dataset (red).

To better demonstrate that RADAR detect DM sites with better sensitivity and specificity in real data, we show examples of DM site that is only detected by RADAR as well as likely false discovery sites identified by exomePeak and Fisher’s test but not by RADAR in the T2D dataset. We plot sequence coverage of individual samples for the DM sites in the RNF213 gene (**Additional file 1: Fig. S7A**) and show despite of large variability in control samples, m^6^A enrichment of T2D samples are consistently lower on this locus. Conversely, in the bogus DM sites detected by alternative methods (**Additional file 1: Fig. S7B, C**), enrichment differences are mainly driven by one or two outlier samples in one group.

To further demonstrate the advantage of using gene-level read counts over local read counts to account for RNA expression level, we repeated the above analysis using post-IP counts adjusted by the local read counts of INPUT. We showed that in the T2D dataset, gene-level adjustment not only enabled stronger signal detection, but also lowered FDR as we observed that the permutation analysis using local counts adjustment resulted in undesired stronger signals around zero in the p-value histogram (**Additional file 1: Fig. S8**). In the ovarian cancer and the mouse liver datasets, local counts adjustment achieved higher signal detection but at the cost of a higher FDR. This analysis suggested that using gene-level read counts as the estimates of pre-IP RNA expression levels could effectively reduce FDR and lead to more accurate DM loci detections.

Attributed to the robust representation of pre-IP RNA expression level using gene-level read counts, RADAR’s performance is more robust to the sequencing depth of INPUT samples. To demonstrate this, we applied RADAR on data created by sub-sampling the read counts of INPUT samples in the T2D dataset so that the sequencing depth is half of the full dataset (average 17.5 million reads). We compared the DM sites detected in the reduced dataset with the results obtained from the full dataset (**Additional file 1: Fig. S9A**). Using a 10% FDR cutoff, RADAR-detected DM sites in the reduced dataset showed the highest overlap with that in the full dataset. MeTDiff and QNB only had a few overlapping DM sites between the sub-sampled and full dataset. Fisher’s test and exomePeak had slightly fewer overlaps comparing to RADAR but had more false discoveries. We further compared the log fold change (logFC) estimates from reduced and full datasets to check their consistency. As a result, we found reduced sequencing depth had the least impact on the logFC estimated by RADAR while the estimates by others are much less reproducible with a shallower sequencing depth (**Additional file 1: Fig. S9A**).

Unlike earlier pipelines that perform DM tests only on peaks identified from peak calling, RADAR directly tests on all filtered bins and reports DM sites. To check if the DM sites reported by RADAR are consistent with known characteristics of m^6^A, we performed de-novo motif search on these sites and found DM sites detected in ovarian cancer, mouse liver and T2D datasets are enriched for known m^6^A consensus motif (**Additional file 1: Fig. S10A**) [18], suggesting DM sites reported by RADAR are mostly true. We also examined the topological distribution of these DM sites by metagene analysis (**Additional file 1: Fig. S10B**). The distributions in ovarian cancer and mouse liver datasets are consistent with the topological distribution of common m^6^A sites, indicating methylation changes occurred in these two datasets were not spatially biased. Interestingly, DM sites detected in T2D dataset are strongly enriched at 5’UTR, suggesting T2D related m^6^A alteration are more likely to occur at 5’ UTR.

### RADAR analyses of m^6^A-seq data connect phenotype with m^6^A-modulated molecular mechanisms

Finally, we investigated whether DM test results obtained from RADAR would lead to better downstream interpretation. In the ovarian cancer dataset, we performed KEGG pathway enrichment analysis on the differential methylated genes (DMGs) detected by RADAR (**Fig. 5A**). We found the detected DMGs were enriched with molecular markers related to ovarian cancer dissemination [19, 20]. For instance, we identified key regulators of the PI3K (enrichment p-value 7.8×10^−5^) and MAPK pathways (enrichment p-value 1.1×10^−4^), including hypo-methylated PTEN and hyper-methylated BCL2 (**Additional file 1: Fig. S11**). Other notable DMGs include key markers of ovarian cancer such as MUC16 (CA-125) and PAX8, as well as genes that play key roles in ovarian cancer biology such as CCNE1 and MTHFR. Conversely, DMGs detected by MeTDiff were only enriched in three KEGG pathways (**Fig. 5B**), most likely due to its inadequate power. We showed through permutation analysis that exomePeak and Fisher’s test results included a significant portion of false positives and could lead to biased downstream interpretations.

**Figure 5.**
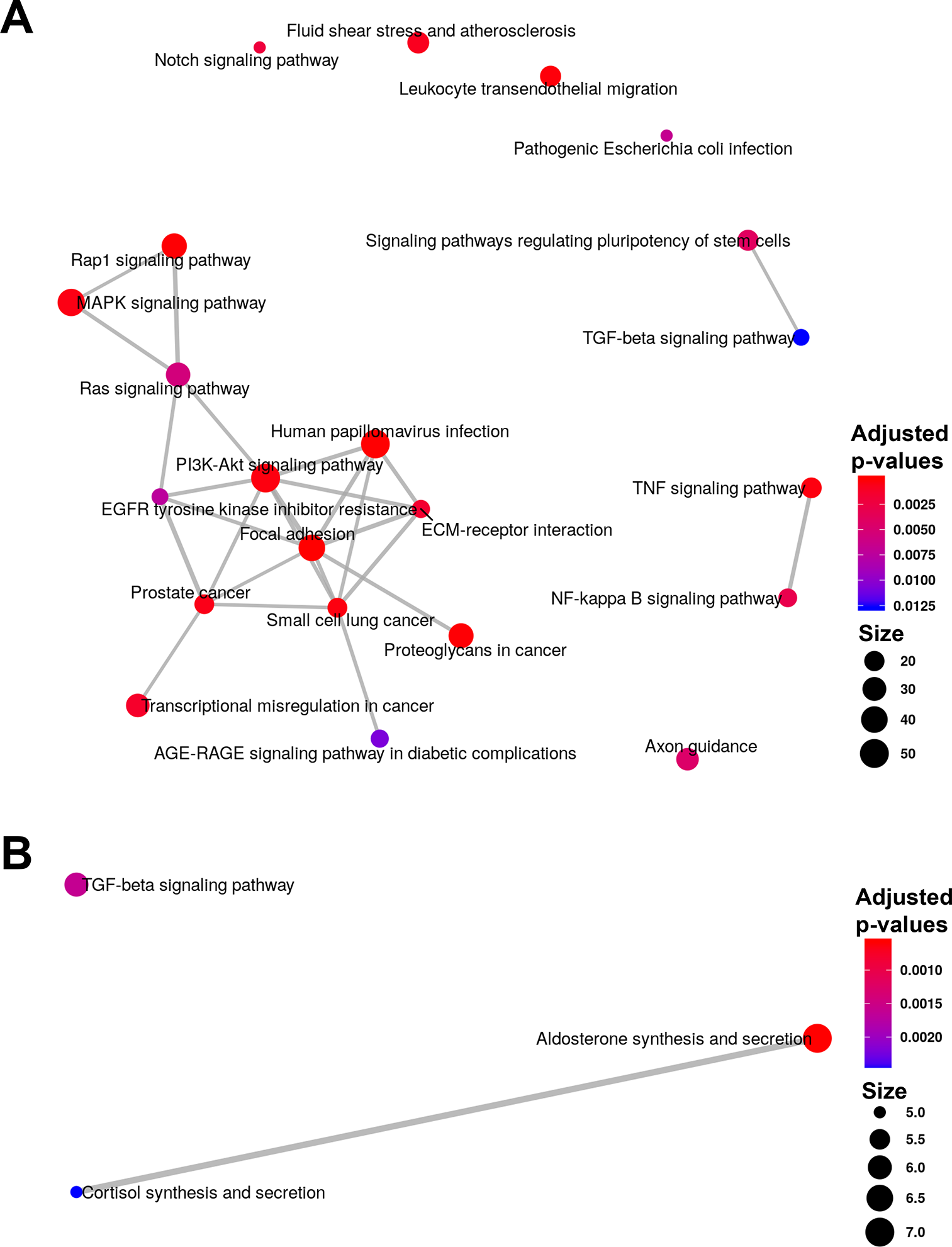
Pathways enriched in differential methylated genes identified in ovarian cancer and T2D datasets. We performed KEGG pathway enrichment analysis using *ClusterProfiler* [37] on DMGs identified in the ovarian cancer dataset by RADAR (**a**) and MeTDiff (**b**), respectively. The enrichment maps represent identified pathways as a network with edges weighted by the ratio of overlapping gene sets.

In the T2D dataset, DMGs identified by RADAR were enriched in related pathways including insulin signaling pathways, type II diabetes mellitus, mTOR pathways and AKT pathways (**Additional file 1: Table S1**), indicating a role that m^6^A might play in T2D. We further analyzed these DMGs in related pathways and found the methylome of insulin/IGF1-AKT-PDX1 signaling pathway been mostly hypomethylated in T2D islets (**Additional file 1: Fig. S12**). Impairment of this pathway resulting in down-regulation of PDX1 has been recognized as a mechanism associated with T2D where PDX1 is a critical gene regulating β-cells identity, cell cycle, and promote insulin secretion [21–24]. Indeed, follow-up experiment on a cell line model validated the role of m^6^A in tuning cell cycle and insulin secretion in β-cells and animal model lacking methyltransferase Mettl14 in β-cells recapitulated key T2D phenotypes (results presented in a separate manuscript, [25]). To summarize, RADAR-identified DMGs enabled us to pursue an in-depth analysis of the role that m^6^A methylation plays in T2D. On the contrary, due to the incapability to take sample acquisition batches as covariates, the alternative methods were underpowered to detect DM sites in T2D dataset and could not lead to any in-depth discovery of m^6^A biology in T2D islets. These examples suggest that MeRIP-seq followed by RADAR analysis could further advance functional studies of RNA modifications.

### Validation of RADAR-detected DM sites by the SELECT method

Recently, Xiao et al. developed an elongation and ligation-based qPCR amplification method (termed SELECT) for single nucleotide-specific detection of m^6^A [26]. This method relies on mechanism different from antibody pulldown-based MeRIP-seq to detect m^6^A, making it a suitable method for validating DM sites discovered by RADAR analysis. We selected 6 DM sites (**Additional file 1: Table S2**) including 2 sites only detected by RADAR and 4 sites in genes important in β-cell for experimental validation using the SELECT method. Among 6 validated sites, the β-cells regulator PDX1 and RADAR-specific DM sites showed significant m^6^A level alteration with p-value 0.009 and 0.017, respectively (**Fig. 6**). Three other sites: IGF1R in the insulin/IGF1-AKT-PDX1 signaling pathway, MAFA—another important regulator of β-cells function and RADAR-specific DM site in CPEB2 showed m^6^A changes consistent with RADAR result despite of not reaching statistical significance. The sites in the TRIB3 gene are similarly methylated in control and T2D samples as measured by SELECT. Overall, five out of six experimentally validated sites were supported by orthogonal evidence by SELECT, confirming the reliability of RADAR-detected differential methylation sites.

**Figure 6.**
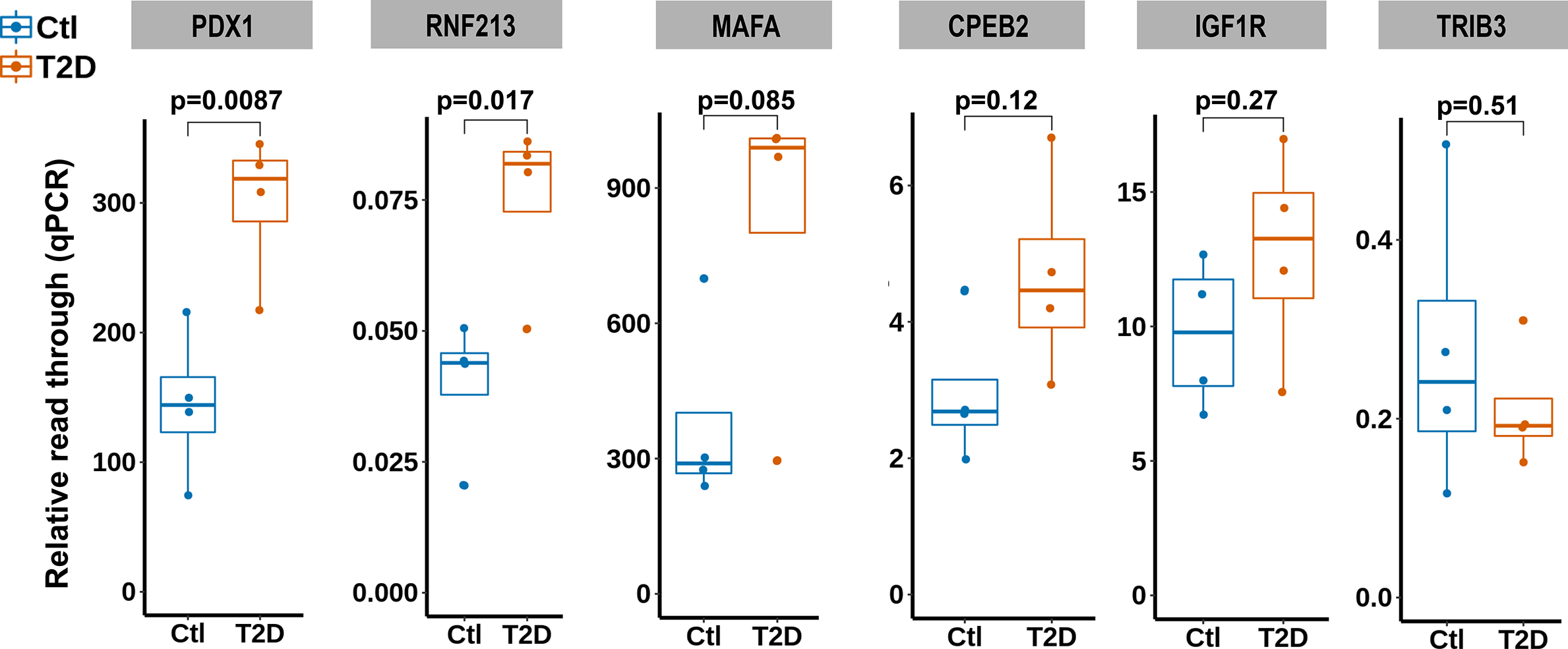
Experimental validation of RADAR-detected DM sites using SELECT method. We applied antibody independent method SELECT on T2D samples (N=4). Shown are SELECT results of 6 putative DM sites for validation. SELECT measures the relative abundance of non-methylated RNA molecules of target locus as represented by the elongation and ligation “read through” of olilgo probes. Thus, SELECT results—“relative read through”—are inversely correlated with m^6^A level.

## Discussion

Sample size is an important parameter of the study design that directly affects the power and reproducibility of an inferential test [27]. DM analyses based on less than 3 biological replicates are still common practice in the RNA epigenetics field and have been shown to exhibit poor reproducibility [28]. To explore the influence of sample size on the power and reproducibility of DM detection, we ran tests on 10 copies of simulated data with effect size equal to 0.75 (roughly two-fold enrichment difference) from two replicates (commonly used) to eight replicates (up-to-date highest). We show that at an FDR cutoff of 10% to select DM loci, FDR increases rapidly as the sample size gets smaller and less than 5 (**Fig. 7**). When the sample size is greater than 6, improvement of FDR is slow while sensitivity climbs rapidly. Our simulation results show the number of replicates greatly influences sensitivity and reproducibility of DM detection as each additional replicate can bring significant gain of area under ROC curve (**Additional file 1: Fig. S13**). To ensure adequate power and reliable DM analysis, we strongly suggest using no less than 5 biological replicates when surveying the alterations in the epitranscriptome of human samples with commonly sequenced library sizes (≥20 million mappable reads).

**Figure 7.**
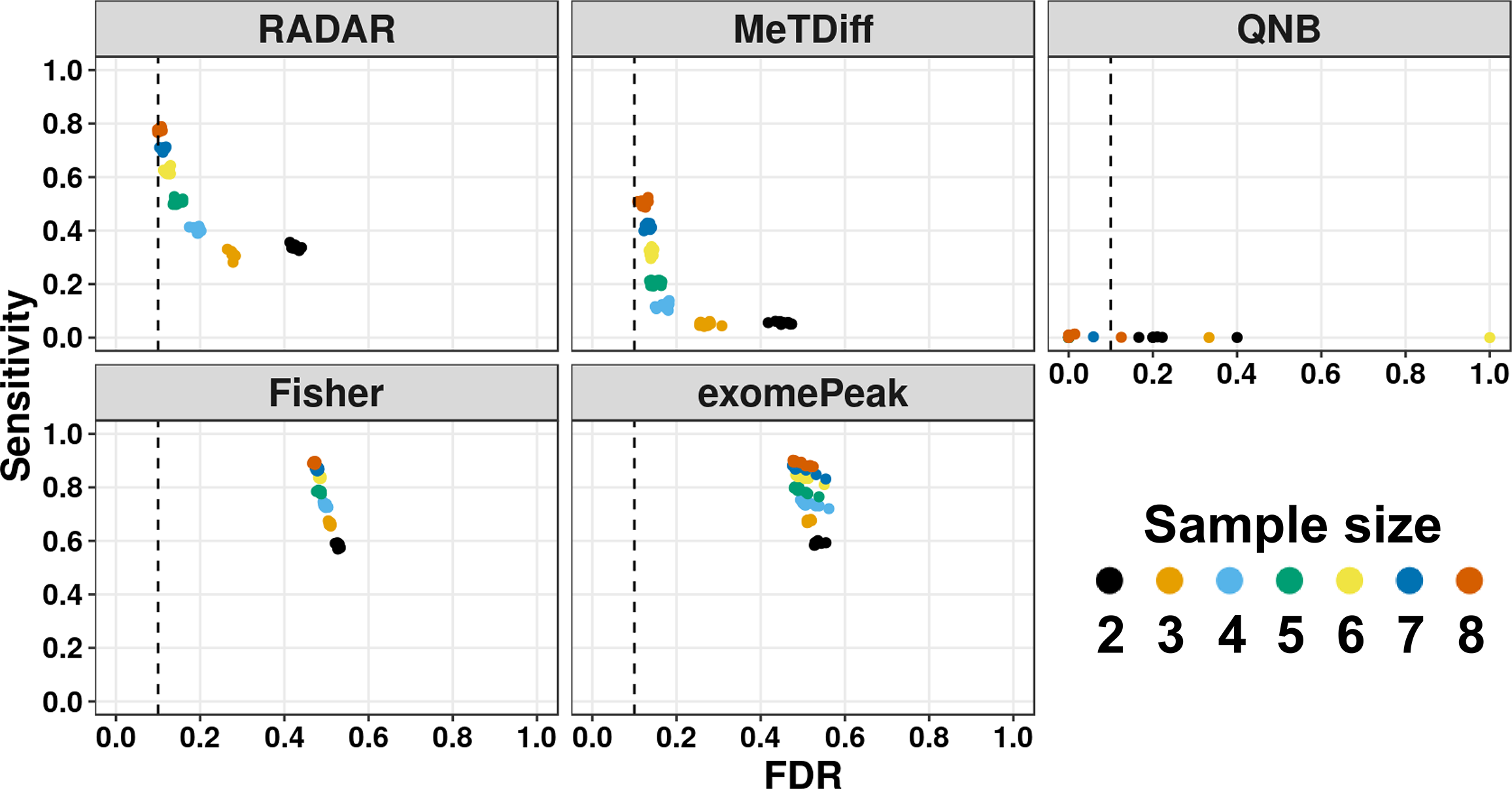
The influence of sample size on the statistical power of differential methylation analysis. Sensitivity vs. empirical FDR for each method on simulated data with different number of replicates (2 to 8) at 10% FDR. Each data point represents the results on one of ten simulated copies. Sample sizes are labeled by colors.

The choice of bin size in the RADAR analysis concerns the tradeoff between resolution and accuracy. At a given sequencing depth, the smaller the bin, the higher resolution achieved. However, a smaller bin size also implies fewer reads in a bin, resulting in increased sampling noise. At common library size of 20 million mappable reads or above, the default bin size (50-bp) should enable sufficient coverage for most enriched regions in the IP library. We recommend to use larger bin size (e.g. 100-bp) for data with smaller library sizes (< 15 million mappable reads) to include enough reads in each bin for DM tests.

A potential caveat of using gene-level instead of local read count to represent pre-IP expression level is that exon-specific expression variation in alternative splicing scenarios would be under-represented. Filtering out DM sites that co-localize with alternative spliced exons in post-processing could help avoid false signals due to this caveat.

## Conclusion

Using simulation and real m^6^A-seq datasets, we demonstrated that RADAR can achieve higher sensitivity with lower FDR than existing methods in the DM analysis. Taking advantage of newly developed SELECT method for experimental validation, we verified that RADAR analysis can uncover true differentially methylated sites. RADAR is a general framework that can be applicable to comparative profiling by MeRIP-seq of various types of RNA modifications including but not limited to *N*^6^-methyladenosine, *N*^1^-methyladenosine and 5-methylcytosine. It also offers great flexibility to adopt to a wide range of mean-variance relationships in the data and accommodate different study designs. We believe RADAR will greatly advance our knowledge of the functions of post-transcriptional modifications.

## Material and Methods

### Ovarian cancer samples

All human tissue samples were collected with informed consent under approved University of Chicago Institutional Review Board protocols and in accordance with the Declaration of Helsinki. All experiments were conducted in accordance with approved protocol guidelines and regulations. Six omental tumor tissues were collected from newly diagnosed patients with advanced, metastatic high-grade serous ovarian cancer during primary debulking surgery at the University of Chicago. Seven normal fallopian tube tissues were collected from patients with benign gynecological conditions at the time of surgery.

### T2D samples

A minimum of 20,000 human islets equivalents (IEQs)/patient were obtained from the Integrated Islet Distribution Program (IIIDP) and Prodo Laboratories. Upon receipt, islets were cultured overnight in Miami Media #1A (Cellgro, USA) and then handpicked, washed twice by self-sedimentation with ice-cold DPBS and pelleted for RNA isolation. All studies and protocols used were approved by the Joslin Diabetes Center’s Committee on Human Studies (CHS#5-05). Samples from eight T2D patients and seven non-diabetic controls were collected for analyses in this study.

### Mouse liver samples

Mouse liver tissues were collected from wild type Albumin-Cre;*Mettl14*^+/+^ and Albumin-Cre;*Mettl14*^flox/flox;Cre^ liver specific conditional knockout mice [29]. Four wide-type and four *Mettl14* cKO mice on 42% high fat diet for three months were sequenced and analyzed.

### RNA extraction and m^6^A-MeRIP-seq

Total RNA was extracted from tissues using TRIzol (Invitrogen) according to the manufacturer’s instruction. For T2D and mouse liver samples, mRNA was further purified with Dynabeads mRNA DIRECT purification kit (Thermo Fisher, cat. 61011). mRNA was adjusted to 15ng/ul in 100ul and fragmented using Bioruptor ultrasonicator (Diagenode) with 30s on/off for 30 cycles. m^6^A-immunoprecipitation (m^6^A-IP) was performed using EpiMark N6-Methyladenosine enrichment kit (NEB cat. E1610S). RNA eluted from m^6^A-IP was cleaned using RNA Clean and Concentrator (Zymo Research, cat. R1013). Input and IP samples were then used to prepare library with KAPA mRNA Hyper Kit (Roche, Cat. KK8541). For fallopian tube and omental tumor tissues, total RNA was fragmented and directly subjected to m^6^A-IP. Takara Pico-Input Strand-Specific Total RNA-seq for Illumina (Takara, Cat. 634413) was used to construct libraries from total RNA where ribosome-derived cDNA was removed before final library amplification. T2D and mouse liver libraries were sequenced by the HiSeq4000 platform at SE50 mode. The ovarian cancer libraries were sequenced by the NextSeq 500 platform at PE37 mode.

### Data preparation

For each dataset, the raw sequencing data were mapped to the corresponding reference genome (hg38 for human, and mm10 for mouse) by *Hisat2* [30] with parameter-x 1. The BAM files obtained from alignment are used as an input file for RADAR.

### Read count pre-processing

RADAR takes a GTF file as an input for gene annotation and obtains a gene model using the R package *GenomicFeatures* [31]. Exons of a gene are concatenated to form the “longest isoform” transcript, which is then divided into bins of user defined size. The R package *Rsamtools* [32] is used to extract and quantify aligned reads from BAM files in each bin. The gene-level counts of INPUT library are obtained by summing up bin-level read counts of each gene.

### Normalization

Unlike previous methods [8–11] that scale read counts to library sizes as a way of normalization, which can be strongly skewed by highly expressed genes [16] (**Additional file 1: Fig. S1**) RADAR considers INPUT and IP samples separately. The INPUT sample is essentially an RNA-seq library; therefore, we directly apply the median-of-ratios method implemented in DESeq2, which is robust to outliers, to estimate a sample-wise size factor for each sample from INPUT gene-level counts. In regard to the IP sample, the abundance of read counts *t*_*i,j*_ depends on the abundance of RNA in the pre-IP RNA pool, the overall IP efficiency of that sample, the total sequencing depth of that IP library, and the methylation level of that locus. To normalize the variation due to sample-wise IP efficiency difference and sequencing depth variations of IP libraries, we estimate a sample-wise size factor from the fold enrichment 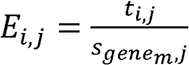 of the top 1% bins ranked by IP read counts where *t*_*i,j*_ is IP read counts and 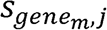 is the normalized *geneSum* (gene-level) read count at corresponding gene. The reason that only top bins are used to estimate the overall IP efficiency is to exclude the regions where IP read counts are mainly attributed to non-specific binding.

### Adjust IP read count for pre-IP RNA expression level

To account for the pre-IP gene expression level variation in IP read counts, we compute a gene-wise size factor by centering normalized gene-level counts to 1. For each bin, we divide the normalized IP read counts by the gene-wise size factor of the corresponding gene. The resulting IP read counts now reflect the methylation level as other factors have all been accounted for. The adjusted IP read counts representing methylation levels are further used for DM tests.

### Data filtering

We apply two filters to remove unwanted bins in the data: 1) We remove bins in which reads are depleted in IP libraries because read counts in these bins are likely attributed to non-specific binding during the immunoprecipitation; 2) We remove bins in which raw IP read counts are smaller than 15 (this cutoff can be defined by the user) because signals in regions without sufficient coverage will be too noisy and unreliable. Low IP read count also implies that the bin is likely a non-methylated region.

### Model for DM test

For each bin, we model the processed IP read counts *Y*_*i*_in the *i*-th sample as follows:

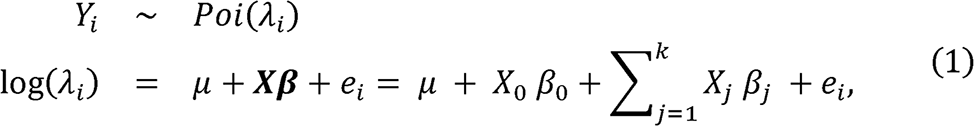

where *λ*_*i*_ is the mean of a Poisson distribution, μ is a bin specific intercept, **X** is the design matrix including the indicator of the groups of interest *X*_0_ and covariates *X*_*j*_ (*j* = 1,…*k*), ***β*** represent associated coefficients and *e*_*i*_ is a random effect following a log gamma distribution with a scale parameter *ψ* and mean equal to 1, i.e., *e*_*i*_ ∈ *logGamma*(*ψ*,*ψ*),. Introducing a new variable *w*_*i*_ ∈ *Gamma*(*ψ*,*ψ*), we have *λ*_*i*_ = *e*^μ+X*β*^*W*_*i*_. The differential analysis is equivalent to test against the null hypothesis *β*_0_ = 0.

After integrating out *W*_*i*_, the likelihood of observing the data given all other parameters *Θ* is:

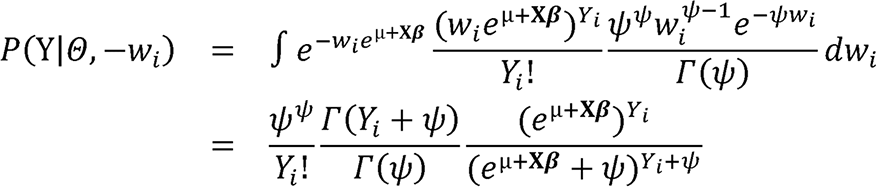

The marginal log likelihood of observing **Y** can be written as:

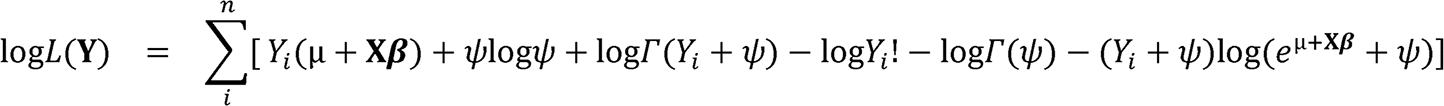

We use the gradient ascent algorithm to calculate maximum likelihood estimators of all the parameters, which involves the calculation of first derivatives.

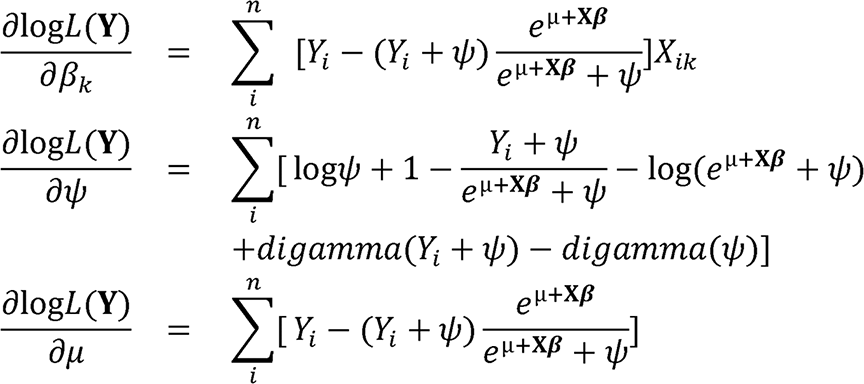

In each iteration, the parameters are updated through 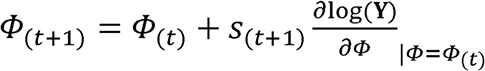. The step size *S*_(*t*+1)_, is determined by a line search algorithm. Finally, a Wald test is derived to test against *β*_0_ = 0, i.e., the test statistics is 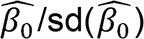 where 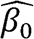 is the MLE and 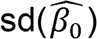 is estimated using observed Fisher information.

### Post-processing

Since the DM tests are performed on consecutive bins on the mRNA, post-processing is needed to merge connected bins that contain reads derived from the same methylation site and report their genome coordinates instead of mRNA coordinates. Specifically, we filter all the bins under user-defined FDR cutoffs and merge adjacent significant bins to a single peak. To represent the mRNA peaks using the genome coordinate, we report the final result in BED12 format, which can specify exon blocks for an intron-spanning interval.

### Simulation analysis

To assess the sensitivity and specificity of each methods on detecting true DM sites, we simulated dataset of 8 replicates with and without covariate. Since RADAR make inferential test on fold enrichment (pre-processed IP read counts adjusted for input RNA level variation), we first simulated this enrichment data (pre-processed IP read counts) using Model (1). We draw the distribution of gene-specific intercept parameter *μ* (equivalent to baseline sequencing depth of control samples), random effect parameter *ψ* from real data (T2D) to better reflect the property of real data. For each dataset, we simulated 26,324 sites where 20% of them were predefined as true DM sites with effect sizes of 0.5, 0.75 or 1. At pre-defined true DM sites, we simulated read counts with ***β*** = 0.5 (or 0.75 or 1) while ***β*** = 0 at null sites. Other alternative methods take INPUT and IP read counts as input data for DM tests. To convert our simulated enrichment data into paired INPUT and IP read counts, we used INPUT read counts from the real T2D dataset and generated corresponding IP read counts by rescaling simulated IP counts to match the IP/INPUT ratio in the real data.

We then applied RADAR, MeTPeak (version 1.1) [8], QNB (version 1.1.11) [10], Fisher’s exact test, exomePeak (version 2.17.0) [11] to the simulated data. We used the Benjamini-Hochberg method to adjust for multiple comparisons. Using an FDR cutoff at 10% (or 1%, 5% … in sliding threshold analysis), we obtained a set of predicted DM sites for each method. We then checked whether pre-defined true DM sites were predicted to be DM site. To evaluate the performance, we computed sensitivity by dividing the number of overlaps between predicted DM sites and true sites by the number of true sites. We also computed the empirical FDR by dividing the number of predicted sites that is not in the true sites by the number of predicted sites. To evaluate effect of covariates on the performance, we repeated the above analysis using Model (1) with an additional binary categorical (such as gender) variable with an effect size of 2. There are 3 covariates included in the T2D data analysis. The covariate with the largest effect size ranges from 0 to 4. We chose an intermediate effect size of 2 to represent a moderately challenging scenario in real data.

Next, we repeated the simulation analysis with an alternative model as described in the QNB manuscript [10], termed as the “QNB model”. Unlike Model (1) that directly simulates the enrichment as pre-processed IP-read counts, the QNB model simulates the paired INPUT and IP read counts separately, each following a negative binomial distribution. The original QNB model sets equal variance for both INPUT and IP data. However, we observe that in the MeRIP-seq, the IP read counts usually have higher variability than the INPUT read counts due to extra variation introduced during the IP process **(Fig. 1C** and **Additional file 1: Fig. S4**). Therefore, we modified the QNB model so that the variance parameter for IP is a magnitude higher than INPUT. Similarly, we generated data for the simple case as well as the difficult case with one confounding factor using the QNB model and applied each method to test for DM loci.

### Coverage sub-sampling analysis

To demonstrate that the robust measurement of pre-IP RNA level implemented in RADAR improves the robustness to varying INPUT sequencing depth, we used T2D dataset as an example and performed sequencing depth sub-sampling analysis. We used the Sambamba [33] with parameters “view -h -t 20 -s 0.5 -f bam-subsampling-seed=1231” to sub-sample half of the reads from BAM files of INPUT samples. To count the overlap between results from sub-sampled and full data, we first obtained filtered bins that are shared in both datasets, then count the bin if it reached significant threshold in both datasets. To compare the log fold change (logFC) estimates, we plotted the logFC estimated from the sub-sampled data against that estimated from the full data by each method.

### *De novo* motif discovery and metagene analysis

To examine if the putative DM sites detected by RADAR are consistent with known characteristics of m^6^A sites, performed *de novo* motif discovery analysis using the *findMotifs* function of homer2 [34] with parameter “-len 5,6 -rna -p 20 -S 5 -noknown”. A background sequence of randomly sampled peaks on transcriptome was used in the motif analysis.

Topological distribution of putative DM sites were plotted on a metagene using R package Guitar [35].

### Pathway enrichment analysis

A pathway enrichment analysis was performed on the DMGs identified from the ovarian cancer dataset using KEGG pathways [36] using the *enrichKEGG* function in R package *ClusterProfiler* [37].

### Experimental validation by SELECT method

SELECT is an elongation and ligation-based method that can distinguish single m^6^A site from A site [26]. Briefly, we design two oligos flanking the target m^6^A/A site that leave a gap on the m^6^A/A site. Then *Bst* DNA polymerase and SplintR ligase are used to fill the gap where m^6^A hinders the elongation of the complementary oligo and thus prevent the gap to be filled. Finally, qPCR targeting the ligated oligo is used to quantify the abundance of the non-methylated RNA molecules. qPCR quantification targeting a nearby region on the target gene is used to normalize the gene expression variation. Since readout of SELECT method reflects relative abundance of non-methylated molecules, we expect the SELECT result to be inversely correlated to the m^6^A levels.

Since SELECT method involves many steps for each site and is not feasible for high throughput analysis, we selected 6 sites including 2 DM sites that were only detected by RADAR and 4 DM sites that were implicated in T2D biology for experimental validation. We first matched RRACH motif in the putative DM peak and designed complementary oligos of 30 nt flanking the putative m^6^A site. An additional 21 nt sequence at 5’ of the up-probe and 20 nt sequence 3’ of the down-probe were added to the oligos as universal primer sequence. For each DM site, we also designed a pair of primer targeting the gene harboring the DM site (see **Additional file 1: Table S3** for oligo and primer sequences).

We applied the SELECT method to 4 control and T2D samples that have enough RNA material leftover from sequencing experiment. For each sample, 50 ng of total RNA was mixed with 0.8ul up-probe and down-probe oligo (1uM) of each target m^6^A site, 1 ul dTTP (100uM), and 2ul 10X CutSmart buffer (NEB) supplemented with H2O to 17ul total volume. The mix was incubated at a temperature gradient: 90°C for 1 min, 80°C for 1 min, 60°C for 1 min, 50°C for 1 min and then 40°C for 6 min. Subsequently, a 3ul of enzyme mixture containing 0.5ul *Bst* 2.0 DNA polymerase (0.02 U/ul) (NEB M0275S), 0.5 ul SplintR ligase (1U/ul) (NEB M0375S) and 2 ul ATP (5mM) was added in the former mixture to the final volume of 20 ul. The final reaction mixture was incubated at 40°C for 20 min then denatured at 80°C for 20 min. Then, qPCR reaction to quantify the “read through” oligos was assembled by 10ul 2X qPCR master mix, 0.8ul universal primer as designed in the oligo probes (10uM), 2ul reaction from previous step and 7.2 ul H2O. To quantify the RNA expression level of each gene harboring the m^6^A site, we first prepared the cDNA from 50ng of total RNA using SuperScript VILO Master Mix (Thermo Fisher 11755050). Then 2ul of cDNA were used for qPCR quantification of each gene. Finally, the gene expression level quantification was used to normalize the “read through” oligo probe quantification to obtain “relative read through” level for each site. Note the “relative read through” level reflect the non-methylated level, which is inversely correlated with m^6^A site.

### Sample size analysis

To investigate the effect of sample size on the power of detecting DM sites, we simulated datasets of varying sample sizes from N = 2 to N = 8 using Model (1) without covariates. We assessed the performance by plotting the sensitivity of each method against its FDR using DM loci obtained at an FDR cutoff of 10%. Additionally, we also made the ROC curve for each sample size by varying the FDR cutoffs when selecting predicted sites (**Fig. S13**).

### Availability of Data and Materials

#### Software Availability

Our method is implemented as a C++/R package and is freely available at: https://github.com/scottzijiezhang/RADAR under GNU General Public License (GPL-v3.0). Reproducible documents with all the analysis presented in the paper are available at: https://scottzijiezhang.github.io/RADARmanual/Reproducible.html.

Source code of the software can be found at Zenodo repository: https://doi.org/10.5281/zenodo.3561035

#### Data Availability

All MeRIP-seq datasets sequenced and analyzed in this manuscript have been deposited in GEO repository: GSE 119168 (ovarian cancer), GSE 120024 (T2D) and GSE 119490 (mice liver). The mouse brain dataset was downloaded from GSE 113781 [7].

## Ethics declarations

### Competing interests

The authors declare that they have no competing interests.

### Ethics approval

Ethics approval is not applicable for this study.

### Authors’ contributions

MC and ZZ conceived the idea, developed the method, and designed the experiments. ZZ implemented the software and performed the analyses. QZ contributed to the simulation model. ZZ, ME, AC, DFDJ and DR designed and performed the experiments (wet). CH, EL, and RNK designed and supervised the experiments (wet). MC, ZZ and CH wrote the paper. All the authors have reviewed, commented and edited the manuscript.

### Funding

MC’s work was supported by the National Institutes of Health (NIH) grants R01GM126553, Alfred Sloan Fellowship and a human cell atlas seed network grant from Chan Zuckerberg Initiative. CH’s work was supported by the NIH RM1 HG008935 and Howard Hughes Medical Institute. RNK’s work was supported by NIH R01 DK67536 and UC4 DK116278.

## Supplementary information

**Additional file 1. Additional figures and tables referenced in the article**.

**Additional file 2. Review history**.

