## Additional file 1 for "RADAR: Differential analysis of MeRIP-seq data with a random effect model"

**Fig. S1**

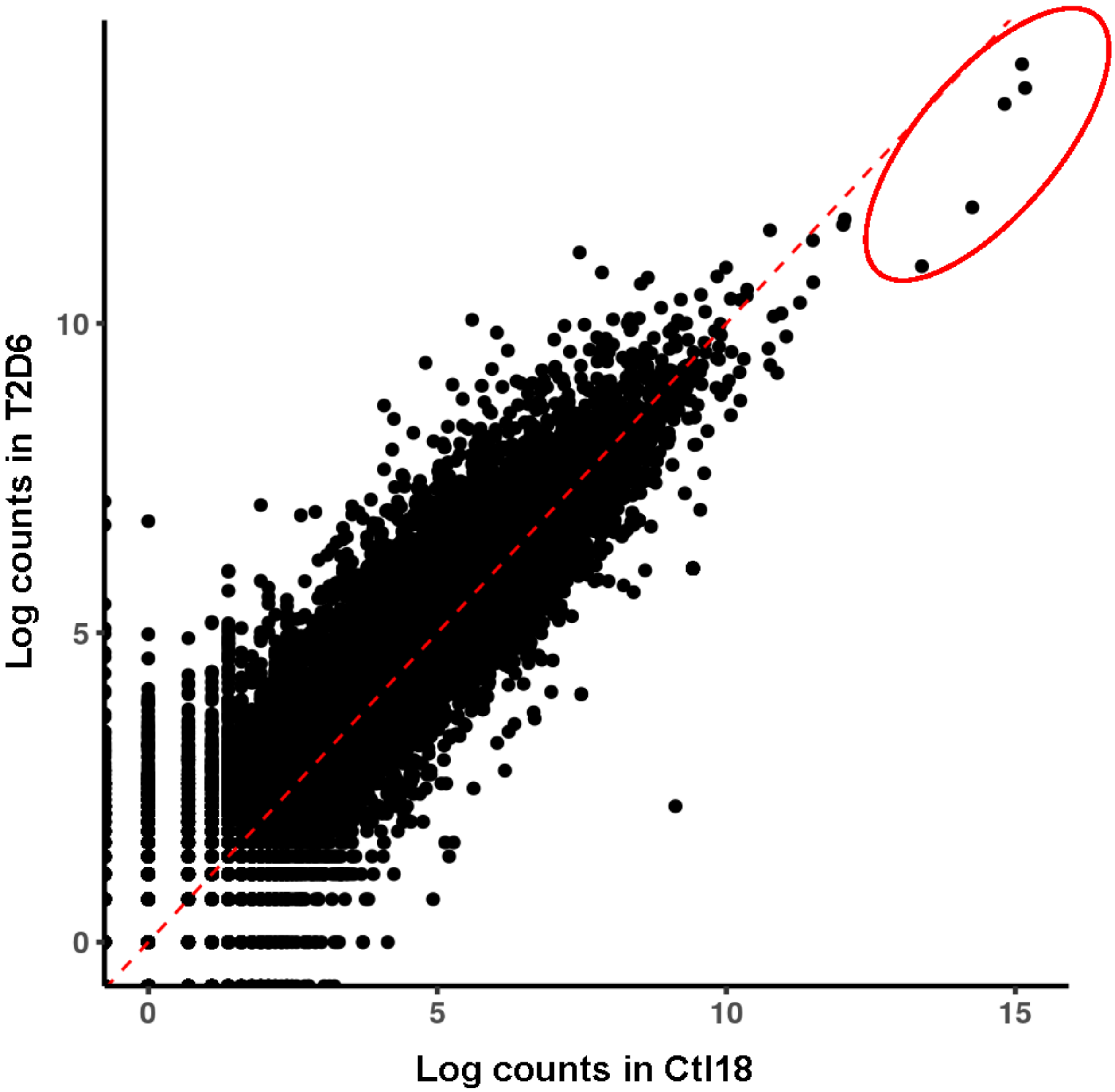


**Fig. S1. Scatter plot of read count.** The input library log read counts of a sample from one experimental group are plotted against a sample from another experimental group. Shown is an example scenario where highly expressed genes can result in underestimation of other genes when normalizing by total coverage. The highly expressed genes that can strongly influence the scaling factor estimation are highlighted by red circles.

**Fig. S2**

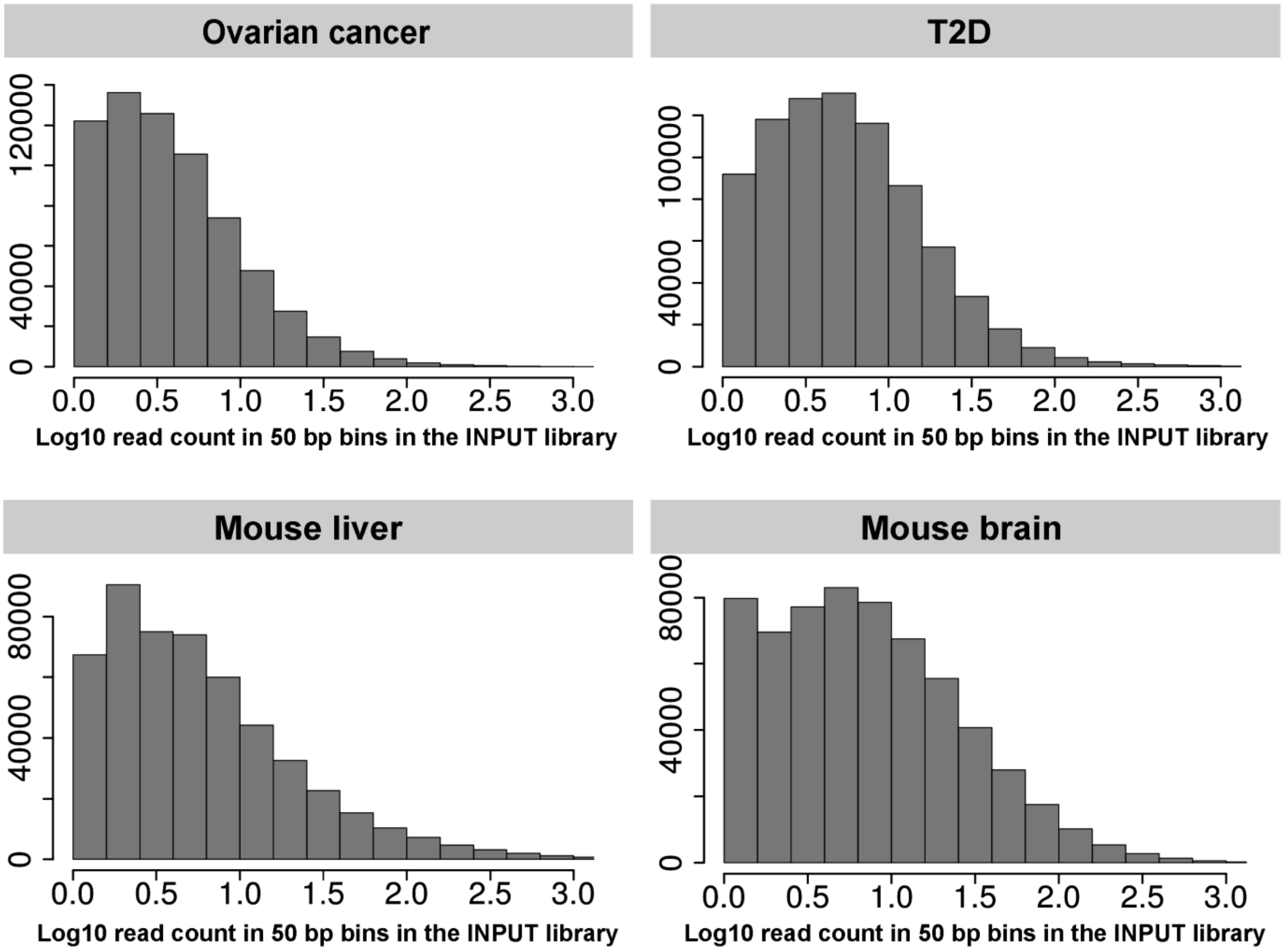


**Fig. S2. Read count distribution of INPUT data.** Distribution of read count $c_i$ (Fig. 1A) in a 50bp bin of input library from real m6A-seq datasets. The read count is shown in log10 scale.

Fig. S3

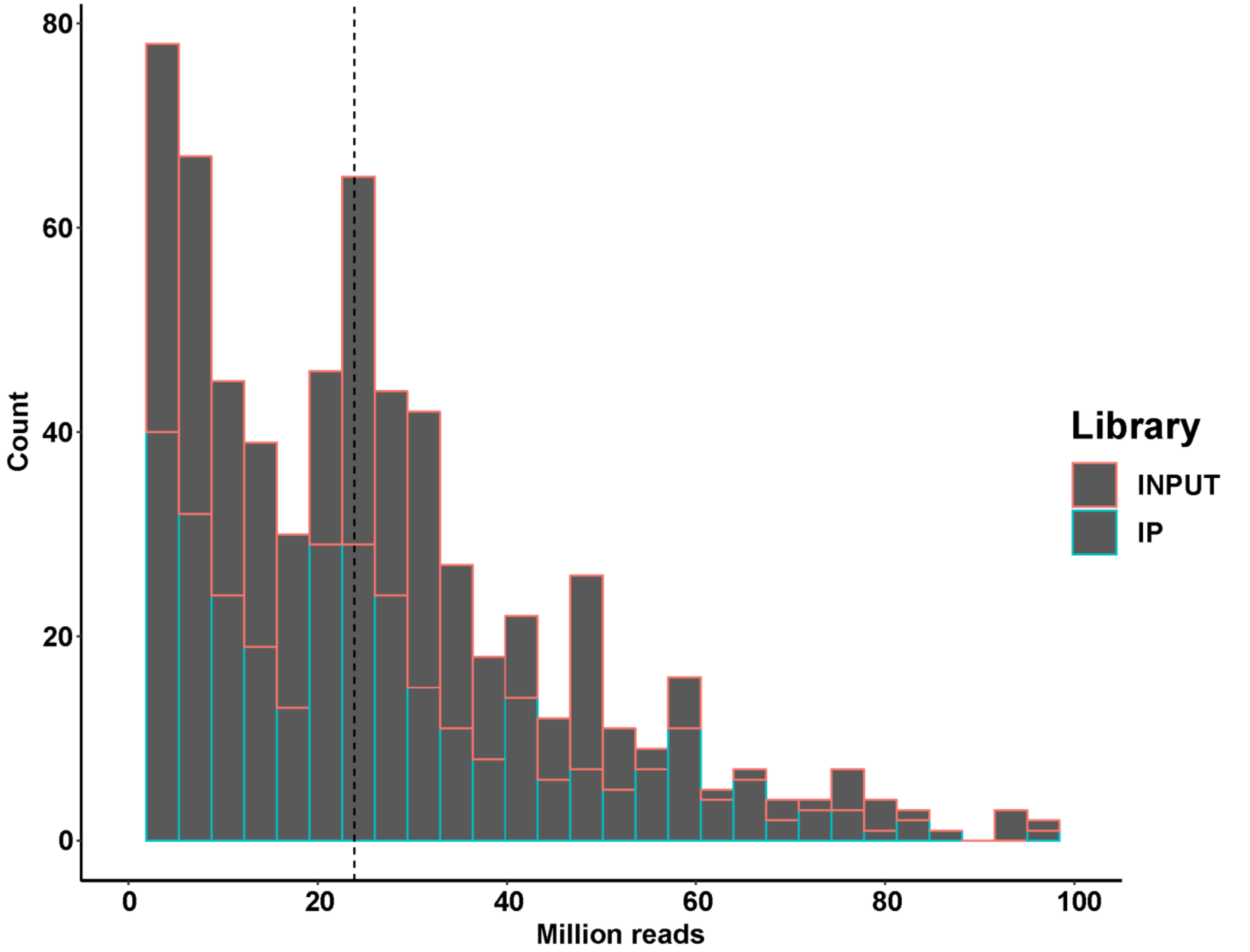


**Fig. S3. Sequencing depth distribution of m6A-seq in published literatures.** Distribution of sequencing depth (million reads) drawn form a m6A-seq database (unpublished observations) is shown by histogram. The database included 339 datasets from published literatures.

**Fig. S4**

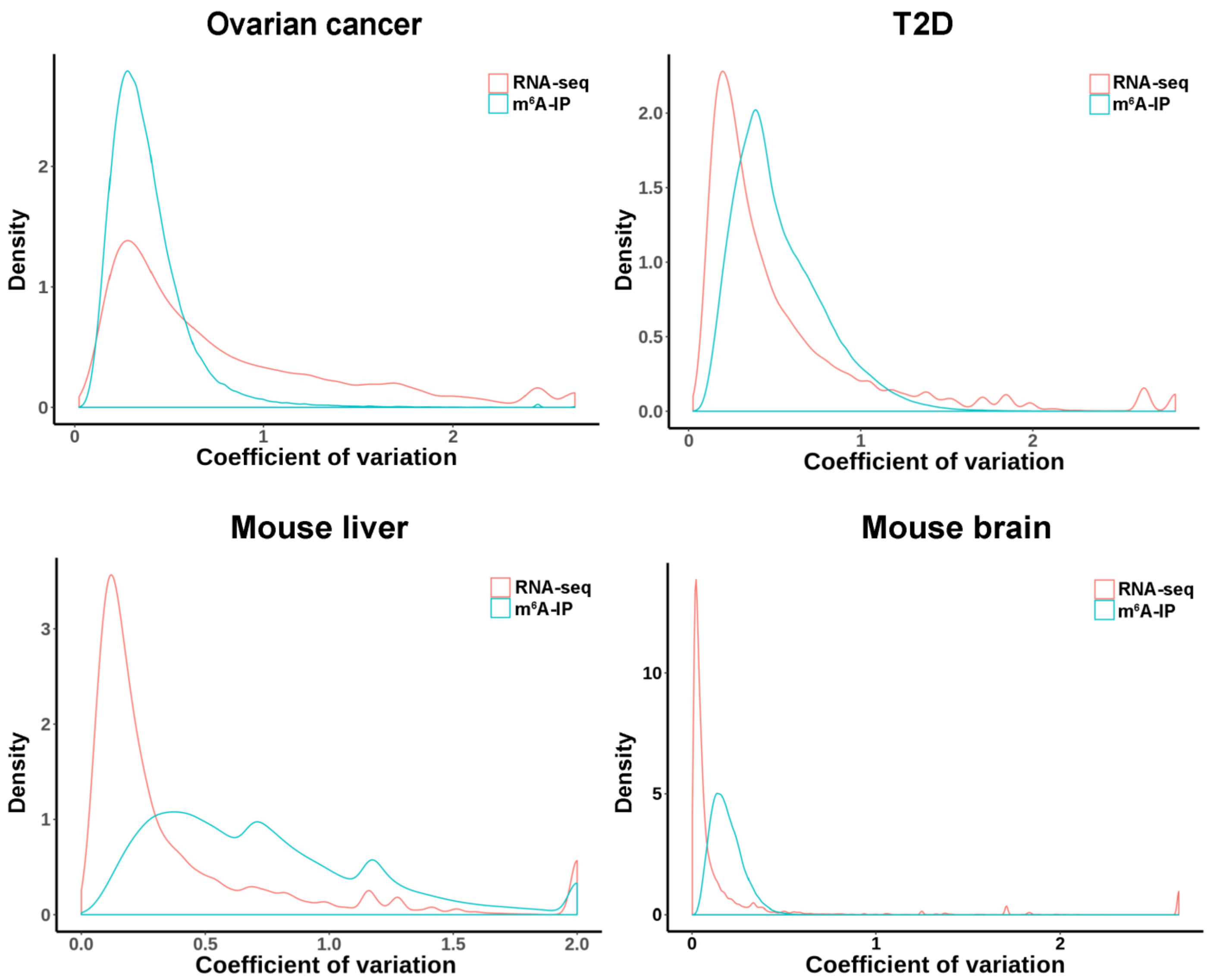


**Fig. S4. Variability distribution comparing RNA-seq and m6A-seq (MeRIP-seq).** Density plot comparing variabilities of m6A-seq data with RNA-seq data. Variability is represented by coefficient of variation.

Fig. S5

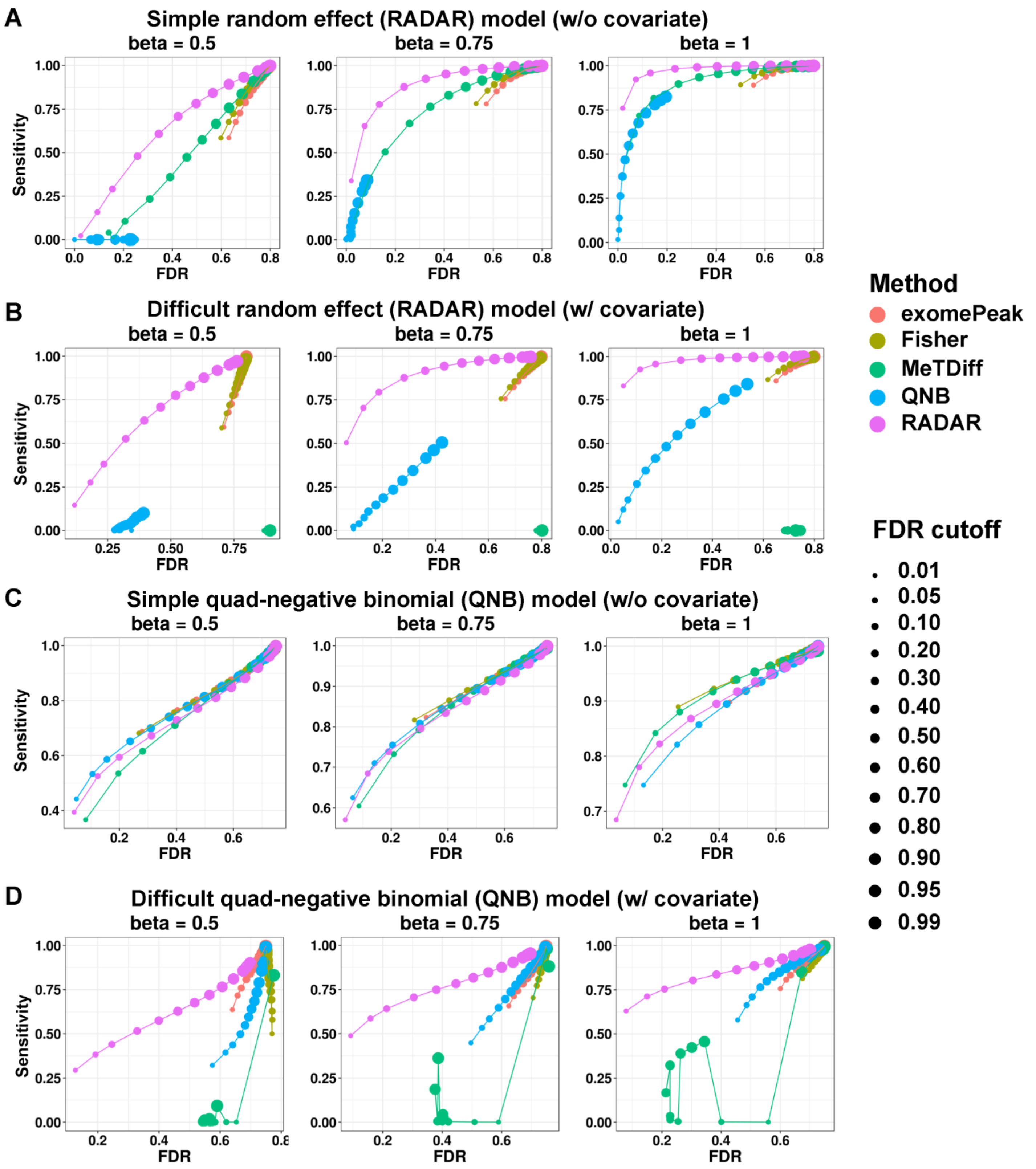


**Fig. S5. Evaluating performances of benchmarked methods on simulated data using sliding thresholds.** We evaluated the performance of RADAR and other methods by comparing the sensitivity and empirical FDR obtained by varying FDR threshold for selecting DM sites. The threshold of selecting DM sites are labeled by the size of data points.

**Fig. S6**

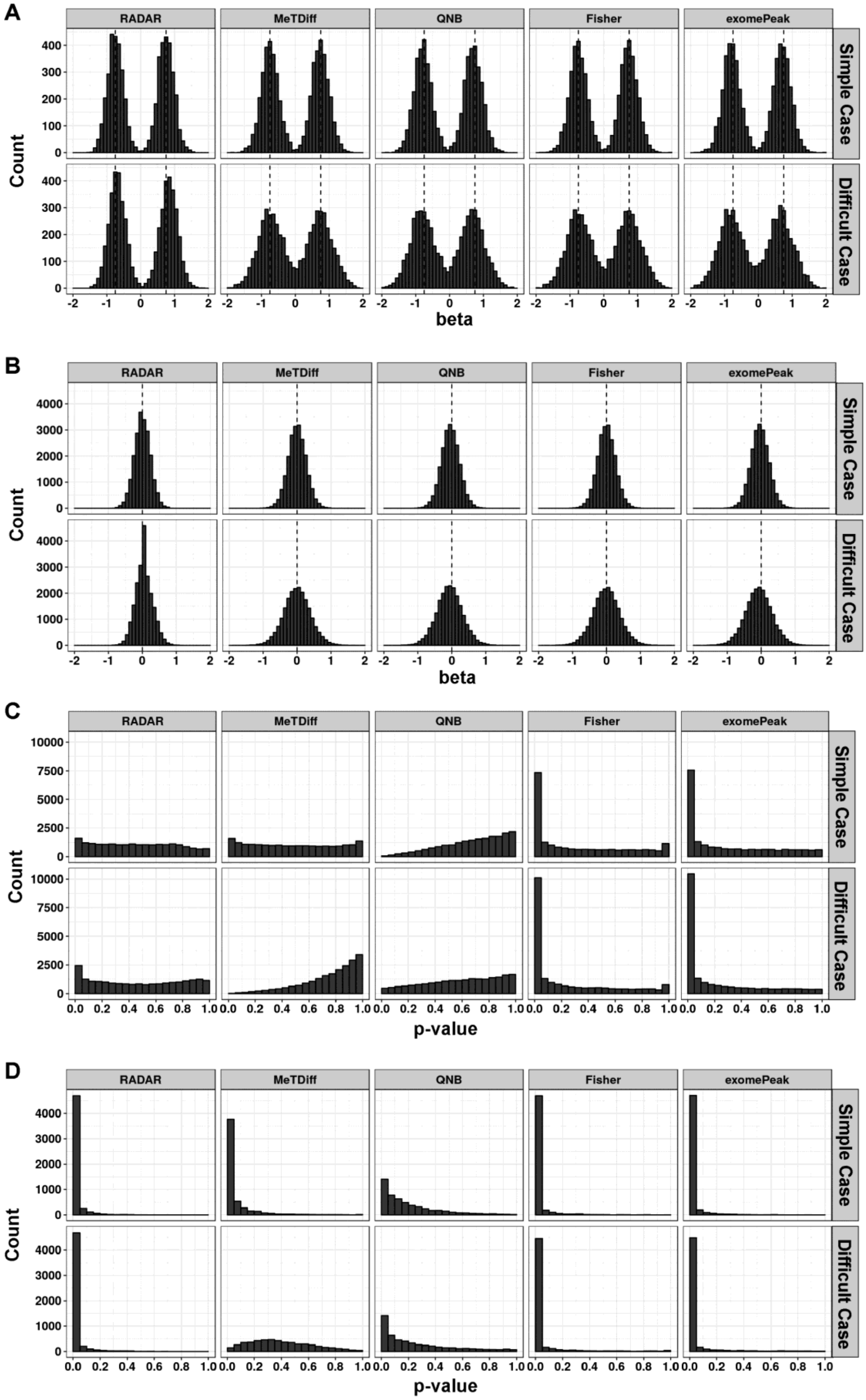
A
B
C
D
RADAR
MeTDiff
QNB
Fisher
exomePeak
Simple Case
Difficult Case
Count
beta
p-value

**Fig. S6 – continued**

**Fig. S6. P-value and effect size estimates on simulated data.** Using data simulated by random effect model of effect size = 0.75, we evaluated the precision of effect size and p-value estimates. (**a**) shows the distribution of effect size estimates in true differential sites and (**b**) shows the distribution of effect size estimates in Null sites where the true effect size is labeled by dashed line. (**c**) shows p-values distribution for Null sites where p-values are expected to be uniformly distributed. (**d**) shows p-values distribution for true differential sites where p-values are expected to be distributed near zero. In all panels, “simple case” refers to simulated data without covariates while “difficult case” refers to simulated data with a covariate.

**Fig. S7**

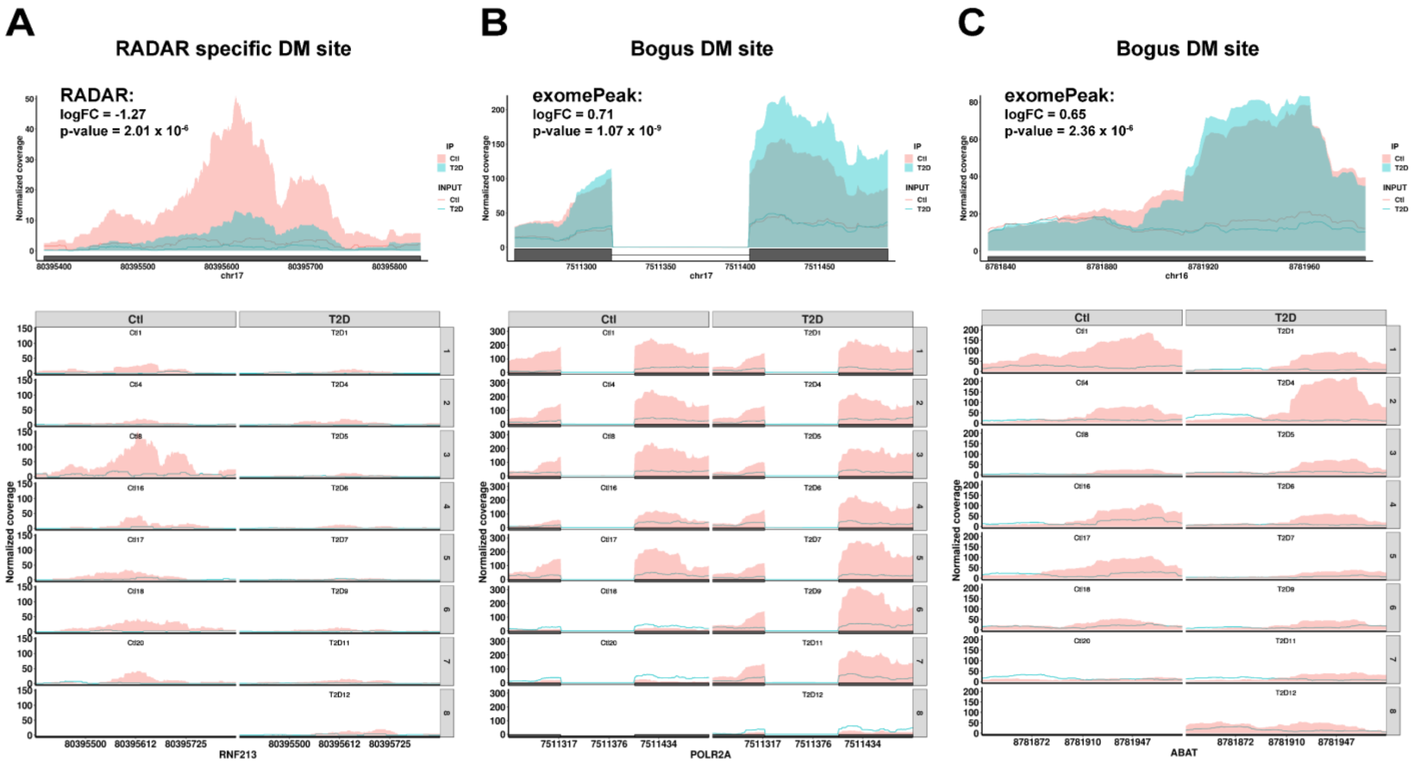


**Fig. S7. Coverage plot of individual samples for example DM sites and bogus sites in the T2D dataset.** We visualize raw data by showing coverage plot for three examples m6A sites. (**a**) shows a putative DM site that was only detected by RADAR but missed by other methods. (**b**) shows a bogus DM site where difference between two groups was mainly driven by two strongly hypomethylated samples in the control group instead of consistent change among replicates. (**c**) shows another bogus DM site that were mainly driven by an outlier hypermethylated sample in the T2D samples.

**Fig. S8**

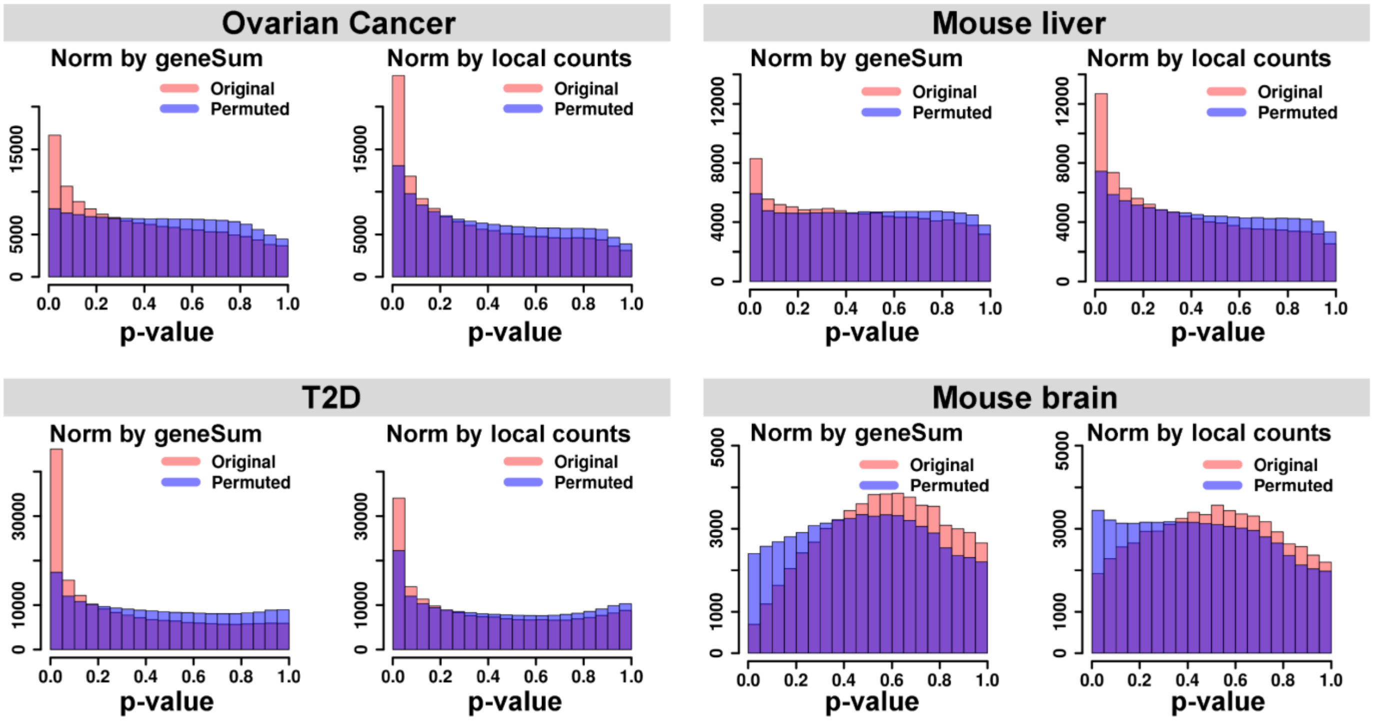


**Fig. S8. Compare the methods to adjust for gene expression level.** Local peak/bin read counts or gene level read counts of INPUT library can be used to account for pre-IP gene expression level variation. We compared the performance of two strategies to adjust for gene expression variation and showed the histogram of p-values from original tests and permutation tests.

**Fig. S9**

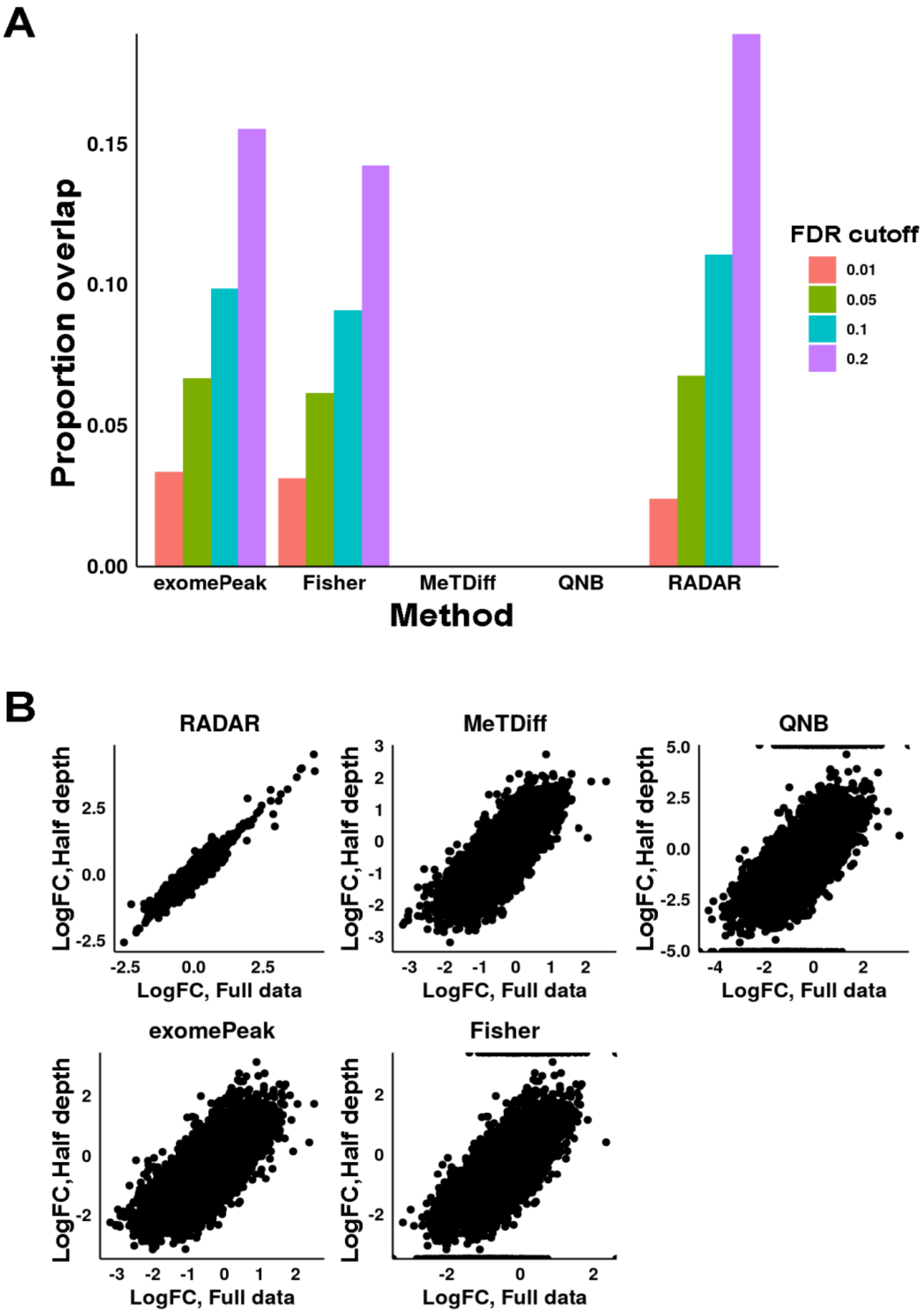


**Fig. S9. Compare results obtained from shallower sequence depth with that from original depth.** We sub-sampled sequence reads from INPUT libraries of the T2D dataset to obtain a dataset of shallower sequence depth (half of the original data). We applied the benchmarked methods to the sub-sampled data and compared the result with the result obtained from the original data. We show the proportion of sites positively identified in both datasets in (**a**) and plotted the estimated log fold changes against each other in (**b**).

**Fig. S10**

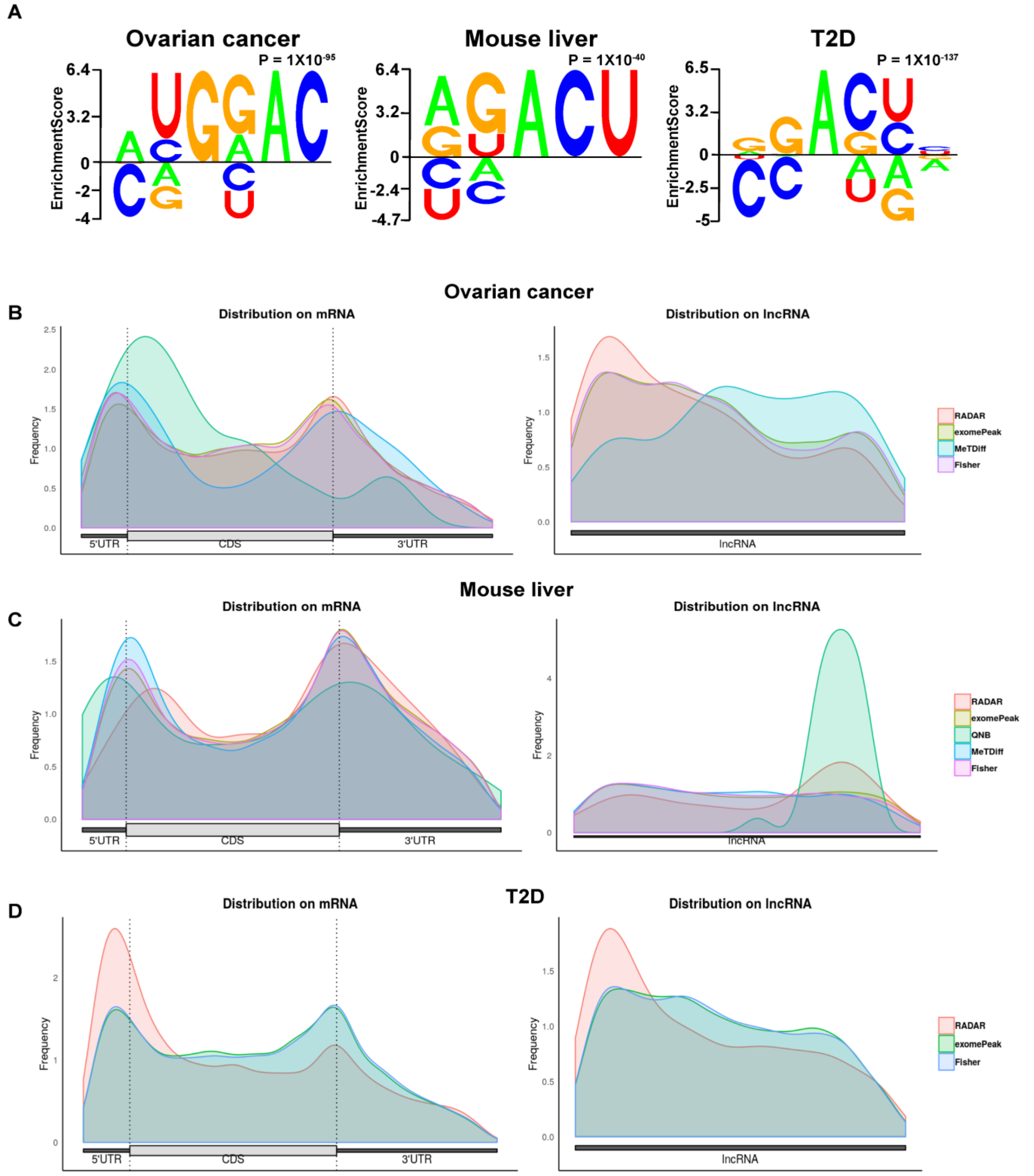


**Fig. S10. Motif analysis and topological distribution of putative DM sites.** We performed de-novo motif search analysis using Homer2 on the putative DM sites detected by RADAR on ovarian cancer, mouse liver and T2D datasets. (**a**) shows RADAR-detected DM sites were enriched for known m6A consensus motif—RRACU. (**b**) shows metagene plots of putative DM sites detected by different methods (method that detected too few DM sites in given dataset were not shown).

Fig. S11

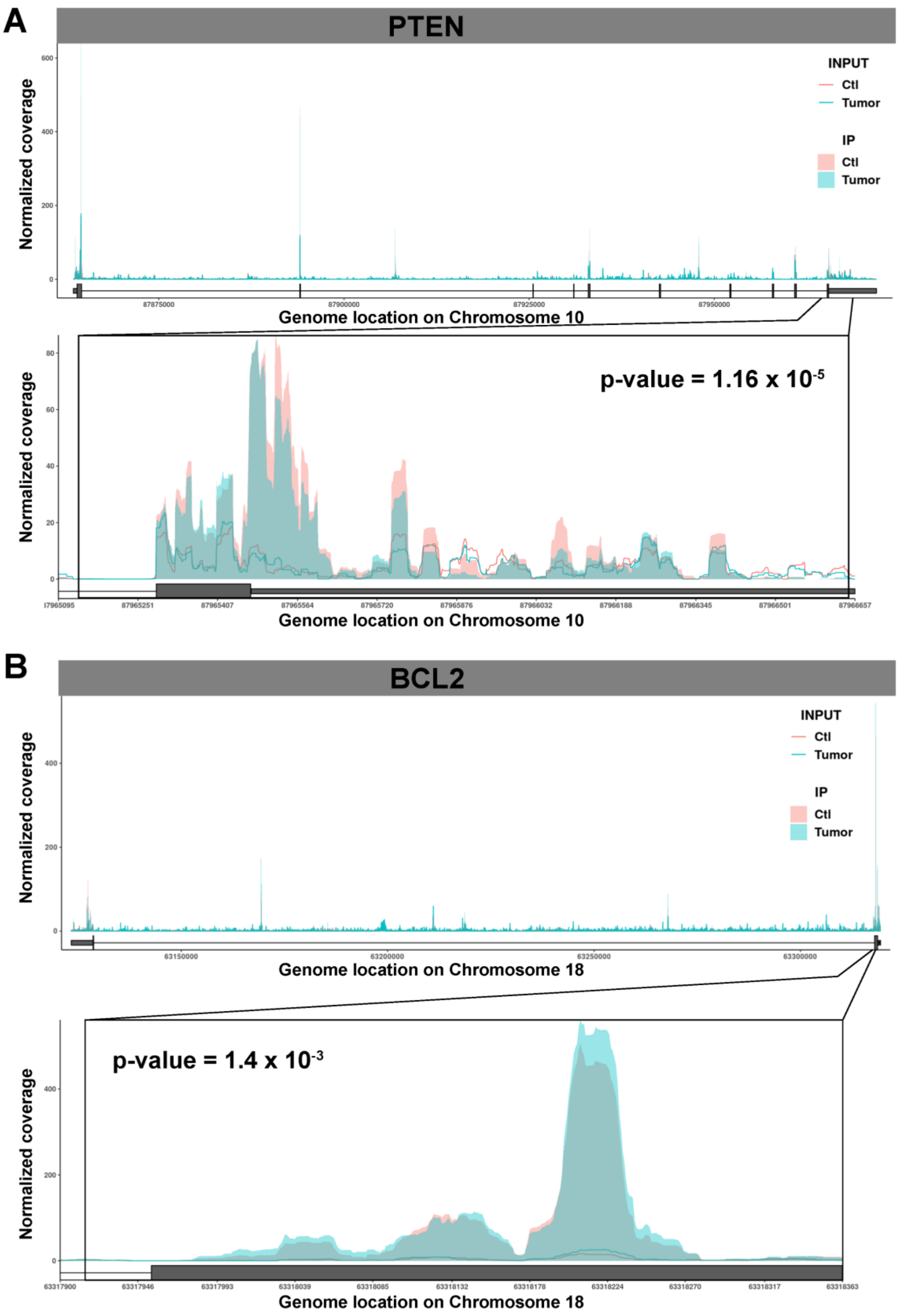


**Fig. S11. Coverage plot to visualize differential m6A peaks in ovarian cancer.** Average coverage of each group is plotted for (**a**) PTEN and (**b**) BCL2. The coverages of both INPUT and IP are normalized by the expression level of target gene so that the coverages of IP samples are directly comparable.

**Fig. S12**

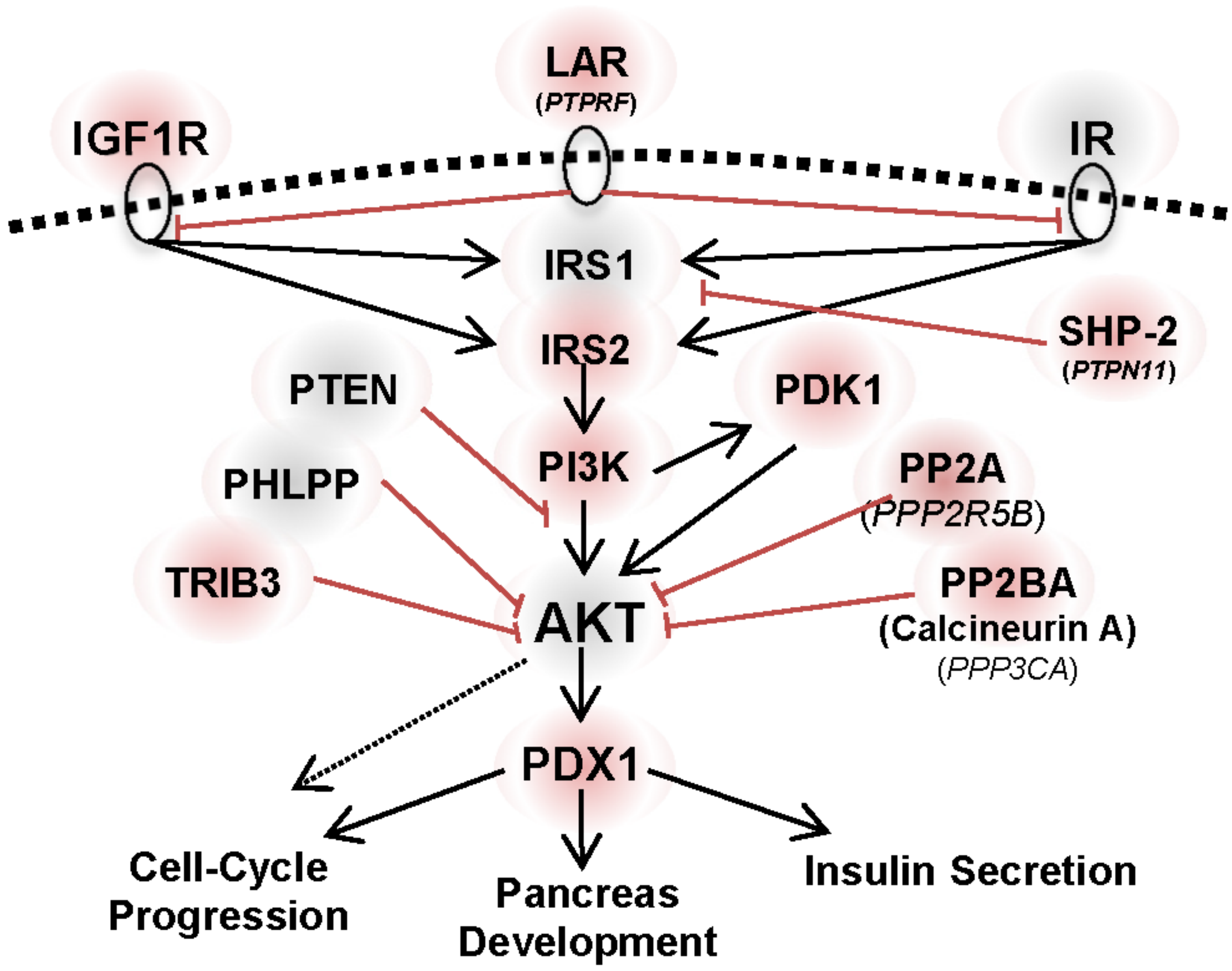


**Fig. S12. Representation of Insulin/IGF1-AKT-PDX1 pathway.** The diagram shows the Insulin/IGF1-AKT-PDX1 signaling pathway based on KEEG and Wikipathway annotations and depicts several $m_6A$ hypomethylated genes (red shade) and unchanged genes (grey shade) in T2D as compared to Controls.

Fig. S13

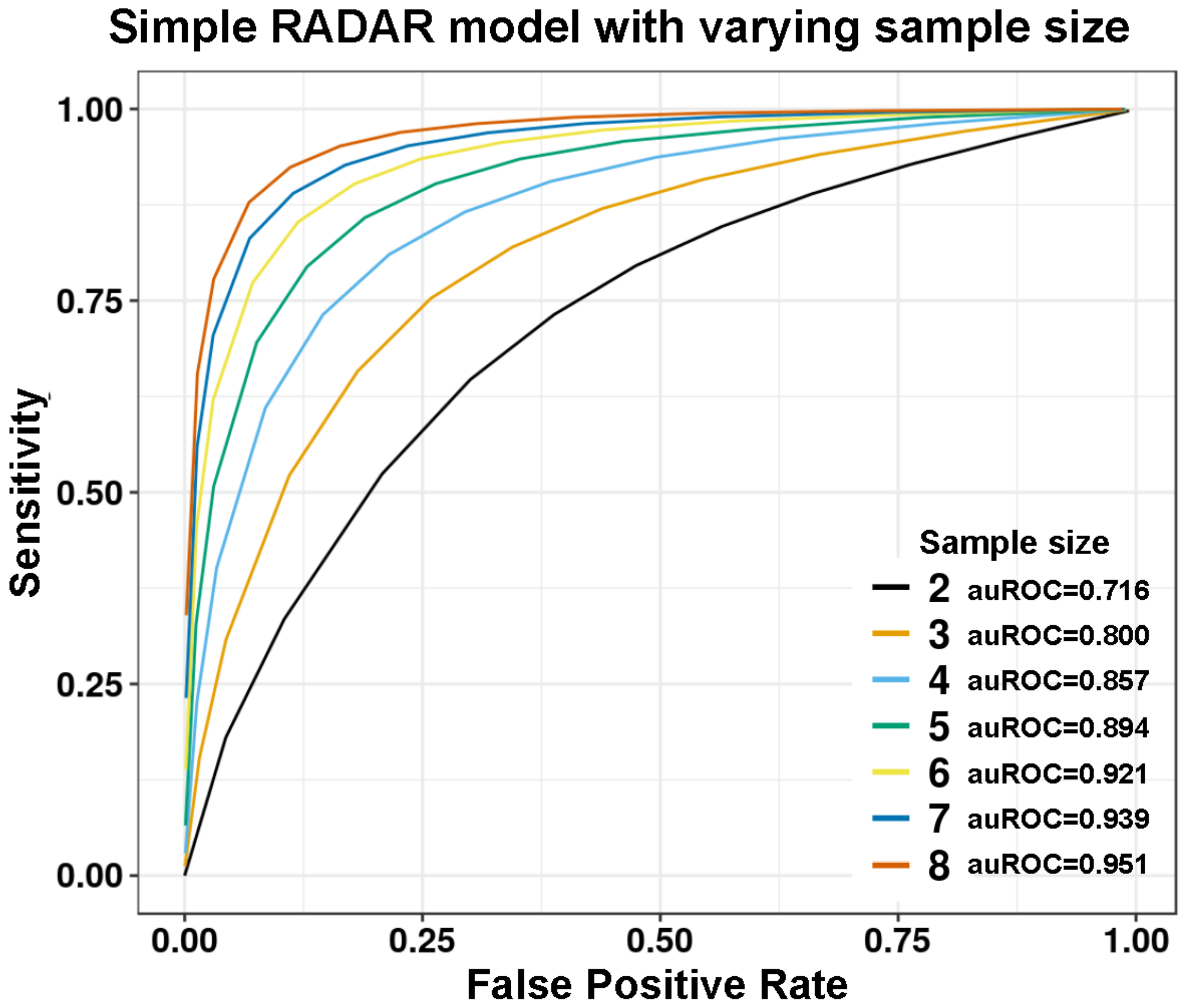


**Fig. S13. Analysis of statistical power and the number of replicates.** We plotted the sensitivity against the empirical FDR by varying the FDR cutoff for selecting predicted differential sites. The larger the area under the curve, the larger the power of the test.

**Table S1**

| Pathway ID | Pathway Names | GeneRatio | BgRatio | pvalue | geneID (Entrez) | Count |
|---|---|---|---|---|---|---|
| hsa04510 | Focal adhesion | 62/1495 | 199/7878 | 2.08E-05 | 3678/3725/2909/1278/9475/2321/29780/10319/7094/3675/3480/3265/55742/7058/3672/5170/2889/10000/3693/394/5595/5500/2534/894/7057/5063/5296/1729/4233/595/1399/5290/2002/858/3910/3918/3915/1292/7422/5156/824/857/896/3694/54776/2932/5602/1284/103910/331/1282/3791/9564/208/2316/5829/2317/3655/5923/6714/81/4659 | 62 |
| hsa04152 | AMPK signaling pathway | 41/1495 | 120/7878 | 5.25E-05 | 5526/6794/8408/1978/1080/3480/23417/5564/5170/10000/5528/84335/57521/2308/5209/2309/9586/8660/90993/5296/31/595/5290/3172/6885/10890/32/5518/51094/5529/3630/5862/5520/7248/5214/23411/6720/10488/208/5525/5527 | 41 |
| hsa04928 | Parathyroid hormone synthesis, secretion and action | 37/1495 | 106/7878 | 7.18E-05 | 4325/3727/1958/4324/2260/4041/2770/5595/9368/9586/9138/6256/5144/2768/2771/90993/3710/11214/5567/2353/10672/846/6667/7421/1026/6257/9365/112/107/2767/4040/369/10488/4205/9935/5332/5566 | 37 |
| hsa04150 | mTOR signaling pathway | 48/1495 | 153/7878 | 0.000144 | 6794/1856/8408/54361/1978/3480/3265/58528/7132/529/5170/10000/4041/84335/57521/79109/5595/64798/8321/55004/54541/96459/5296/5290/79726/90423/64121/9681/1857/83667/54468/84219/1975/3630/10325/23175/7248/4040/2932/3551/9470/389541/208/57600/9894/201163/8140/6520 | 48 |
| hsa03015 | mRNA surveillance pathway | 32/1495 | 91/7878 | 0.000185 | 5526/7919/8106/26528/22985/5528/80335/5500/51585/65109/65110/100529063/23049/8189/9887/79869/53918/5518/5529/10250/5520/10482/140886/8761/5525/23293/5527/22794/1477/53981/10914/2935 | 32 |

| | | | | | | |
|---|---|---|---|---|---|---|
| hsa04933 | AGE-RAGE signaling pathway in diabetic complications | 34/1495 | 100/7878 | 0.000248 | 3725/1278/1958/3265/10000/4089/4088/2308/5595/7046/113026/183/1027/4790/5970/5296/1729/595/5290/7040/7422/6777/581/5590/5602/5580/5581/1284/1282/7042/208/51196/7056/5332 | 34 |
| hsa04068 | FoxO signaling pathway | 42/1495 | 132/7878 | 0.000262 | 100132074/1387/6794/9455/9454/3480/3265/5564/5170/10000/4089/5934/4088/2308/5595/2309/7046/894/2033/1027/8660/5296/595/6789/5290/4193/7040/1026/3630/23411/1017/5602/3551/4303/1901/1454/7042/369/208/1032/1030/7874 | 42 |
| hsa04530 | Tight junction | 51/1495 | 170/7878 | 0.000311 | 776/3725/6794/93643/9475/4771/1080/5962/23370/7122/5564/11346/4628/9368/9414/1740/8189/51421/9075/23327/595/56288/5567/4637/7082/1364/83700/100506658/5518/79784/10096/154796/84952/79778/5590/5520/64398/1365/5602/5581/1741/103910/4214/137075/8976/9223/8777/6714/81/5566/1739 | 51 |
| hsa04010 | MAPK signaling pathway | 80/1495 | 295/7878 | 0.000314 | 3481/775/773/776/3725/6237/1843/3727/4215/6722/2321/9479/2260/3304/2005/3480/3265/1847/7132/8912/9448/100506012/9175/10000/5530/1850/5595/7046/5598/1435/3925/23162/3310/2768/4790/23542/5970/4233/5606/5567/6789/2353/3554/1399/22800/80824/2002/5494/7040/6885/4609/4803/7422/774/3303/5156/3630/7186/5602/781/3551/7042/8605/4137/369/3791/4214/208/3305/2316/51347/4775/7039/9261/2317/23118/9254/5923/57551/5566 | 80 |
| hsa04950 | Maturity onset diabetes of the young | 13/1495 | 26/7878 | 0.000345 | 3651/389692/4821/6928/3087/6927/3170/3171/3172/3110/3630/168620/222546 | 13 |

| | | | | | | |
|---|---|---|---|---|---|---|
| hsa04071 | Sphingolipid signaling pathway | 38/1495 | 119/7878 | 0.000466 | 5526/9475/56848/3265/7132/4363/5170/10000/5528/2770/8877/5595/29956/2534/130367/2768/4790/2771/6609/5970/5296/10672/5290/5518/5529/581/5590/5520/7186/9846/5602/1901/5581/259230/208/5525/5527/5332 | 38 |
| hsa04931 | Insulin resistance | 35/1495 | 108/7878 | 0.000567 | 7132/5564/5170/10000/51085/2308/5500/22877/8473/5781/9586/183/57761/11000/4790/8660/90993/5970/5296/5290/6945/32/3630/5590/5465/2932/5602/5580/3551/5581/6720/10488/208/5836/4792 | 35 |
| hsa04520 | Adherens junction | 25/1495 | 72/7878 | 0.001119 | 1387/9855/2260/10163/3480/5777/4089/4088/5595/117178/7046/2534/2033/4233/56288/7082/10580/6885/8826/4008/999/8976/5797/6714/81 | 25 |
| hsa04930 | Type II diabetes mellitus | 18/1495 | 46/7878 | 0.001147 | 775/773/776/3651/389692/5595/8660/5296/5290/774/3630/5590/5602/5580/80201/3551/5581/122809 | 18 |
| hsa04919 | Thyroid hormone signaling pathway | 36/1495 | 119/7878 | 0.001924 | 1387/9862/10231/54361/7067/3265/5170/10000/488/2308/5595/113026/23389/6256/2033/6567/8202/5296/6548/595/5567/5290/4193/4609/6257/4855/9969/5214/2932/5469/208/51196/4853/5332/6714/5566 | 36 |
| hsa04390 | Hippo signaling pathway | 44/1495 | 154/7878 | 0.002303 | 1856/9113/54361/4771/26524/7003/4089/4088/5500/7532/7046/8321/324/894/1740/595/55233/8313/56288/3689/656/23286/1857/7040/4609/7529/4092/5518/154796/8994/896/5590/5520/64398/2932/10413/999/1741/1454/7042/3398/84962/1490/1739 | 44 |
| hsa03040 | Spliceosome | 38/1495 | 135/7878 | 0.005799 | 24148/7919/6625/1659/9785/3304/23020/10291/51362/23451/22985/84991/10929/494115/51729/3310/10713/988/10946/6100/9343/3303/343069/1655/9092/5093/4670/8175/10084/100534599/11325/57461/3305/144983/23350/58517 | 38 |

| | | | | | | |
|---|---|---|---|---|---|---|
| | | | | | /9879/26121 | |
| hsa04210 | Apoptosis | 38/1495 | 136/7878 | 0.006614 | 2021/3725/3265/2081/7132/8837/5170/10000/823/153090/5595/1514/142/4790/3710/5970/5296/2353/8739/5290/1522/6709/4803/824/581/7186/5366/5602/3551/4170/331/143/208/1512/1519/4792/598/8772 | 38 |
| hsa04066 | HIF-1 signaling pathway | 31/1495 | 109/7878 | 0.010223 | 4055/1387/2321/112399/7037/1978/817/3480/10000/5209/5595/2033/1027/4790/5970/5296/5290/5163/7422/3939/1026/3630/7076/5214/229/3162/80201/9470/112398/208/54583 | 31 |
| hsa04392 | Hippo signaling pathway - multiple species | 11/1495 | 29/7878 | 0.013318 | 8642/9113/4771/26524/7003/55233/23286/8994/10413/1454/84962 | 11 |
| hsa04024 | cAMP signaling pathway | 54/1495 | 214/7878 | 0.013347 | 775/776/1387/3725/6237/5443/9475/817/1080/5727/10000/2770/488/5595/5500/9586/2696/116/2149/2033/5139/5144/4790/2771/90993/5970/5296/6548/1908/5567/2353/2740/5290/22800/6662/6752/84152/112/135/64411/1909/2891/5465/107/5602/10488/2693/208/6751/51196/153/4792/5566/4659 | 54 |
| hsa04151 | PI3K-Akt signaling pathway | 84/1495 | 354/7878 | 0.013348 | 3481/5526/3678/6794/1278/2321/2260/1978/3566/10319/3675/3480/3265/3326/2791/118788/7058/3672/5170/10000/5528/3693/5934/57521/5595/5585/2309/7532/9586/894/1435/54541/6256/2149/7057/1027/4790/90993/9170/5970/5296/4233/595/5290/4193/3910/3918/3915/1292/4609/4803/7422/7529/1026/5518/7184/4515/1975/5529/5156/3630/896/3694/5520/7248/1017/2932/3551/9470/4170/1284/5586/1282/3791/10488/208/5525/9223/5527/2790/7039/23239/598/3655 | 84 |

| | | | | | | |
|---|---|---|---|---|---|---|
| hsa04142 | Lysosome | 33/1495 | 123/7878 | 0.019934 | 4669/967/1203/8218/3482/4864/3425/1514/9583/8120/2990/6609/23163/10312/1522/6448/79158/410/2548/8907/8943/256471/55353/84572/1212/10947/162/8546/1512/3074/1519/23431/23062 | 33 |
| hsa04120 | Ubiquitin mediated proteolysis | 36/1495 | 137/7878 | 0.021349 | 55585/63893/7327/22954/23295/27338/10075/23221/9320/51343/64750/9690/54926/89910/8451/23327/57448/51366/23759/4193/7322/5371/997/9246/7321/65264/8924/331/7326/4214/92912/11060/55958/4281/9616/51433 | 36 |
| hsa04910 | Insulin signaling pathway | 36/1495 | 137/7878 | 0.021349 | 1978/3265/5261/5564/5170/2889/10000/57521/2308/5595/5500/8660/5296/31/5567/1399/5290/10580/2002/32/3630/5576/5590/7248/3636/2932/5602/80201/3551/9470/6720/122809/369/208/5836/5566 | 36 |
| hsa04310 | Wnt signaling pathway | 41/1495 | 160/7878 | 0.022279 | 1387/3725/5176/1856/9475/54361/817/59343/1487/4041/5530/4089/4088/6425/8321/324/894/23002/2033/595/8313/5567/166336/1857/4920/6885/4609/5467/6423/80319/896/27130/4040/2932/5602/1454/8061/85409/4775/5332/5566 | 41 |
| hsa01522 | Endocrine resistance | 27/1495 | 98/7878 | 0.023827 | 3725/3480/3265/10000/1031/5595/1027/8202/5296/595/5567/2353/5290/4193/6667/1026/4855/112/581/107/5602/369/5469/208/4853/6714/5566 | 27 |
| hsa04330 | Notch signaling pathway | 15/1495 | 48/7878 | 0.028337 | 1387/9612/1856/1487/2033/55534/1840/113878/1857/4855/5664/6868/9794/151636/4853 | 15 |
| hsa05202 | Transcriptional misregulation in cancer | 46/1495 | 186/7878 | 0.02959 | 84444/2321/2005/3480/466/51274/905/5914/1031/2130/6935/2308/4297/1025/6692/8148/894/3087/6256/1027/4790/221037/5970/9915/4233/4299/4221/4193/5546/904/6667/4609/4094/1026/6257/5371/8938/581/8861/1655/5327/33 | 46 |

| | | | | | | |
|---|---|---|---|---|---|---|
| | | | | | 98/3486/648/598/6929 | |
| hsa04927 | Cortisol synthesis and secretion | 19/1495 | 65/7878 | 0.029637 | 775/776/5443/8912/3777/5151/9586/183/90993/3710/5567/6667/112/107/2767/949/10488/5332/5566 | 19 |
| hsa04371 | Apelin signaling pathway | 35/1495 | 137/7878 | 0.0342 | 6237/1958/56848/3265/4899/2791/5564/10000/4089/2770/8877/4088/5595/7046/30849/2771/3710/6548/595/5567/10672/22800/5289/112/6543/107/5327/999/5581/208/1490/4205/2790/5332/5566 | 35 |
| hsa04722 | Neurotrophin signaling pathway | 31/1495 | 119/7878 | 0.034682 | 3725/4215/817/3265/5170/2889/10000/5595/2309/5781/5598/4790/5970/5296/1399/5290/4803/10782/5664/581/25/10818/2932/5602/5580/3551/4214/208/25970/9261/4792 | 31 |
| hsa04512 | ECM-receptor interaction | 24/1495 | 88/7878 | 0.035636 | 3678/1278/2812/10319/3675/9899/1605/7058/3672/3693/960/9900/7057/3910/3918/3915/1292/6382/255743/3694/1284/1282/375790/3655 | 24 |
| hsa04611 | Platelet activation | 32/1495 | 124/7878 | 0.036476 | 2909/1278/9475/2812/7094/10000/2770/5595/5500/2534/23365/9138/2149/2771/3710/5296/5567/10672/9002/5290/6786/112/5590/107/5739/103910/8605/208/5332/6714/5566/4659 | 32 |
| hsa04915 | Estrogen signaling pathway | 35/1495 | 138/7878 | 0.037743 | 3725/5443/3304/3265/3326/2775/10000/5914/2770/5595/9586/3310/2771/90993/3710/8202/5296/5567/2289/2353/5290/6667/7184/3303/112/107/5580/25984/10488/208/3305/7039/5332/6714/5566 | 35 |
| hsa04657 | IL-17 signaling pathway | 25/1495 | 93/7878 | 0.038246 | 3725/3727/3934/3326/5595/5598/9618/4790/23765/5970/2353/6885/7184/4312/7186/2932/5602/3551/8061/7128/5596/51433/4792/23118/8772 | 25 |
| hsa05418 | Fluid shear stress and atheroscler | 35/1495 | 139/7878 | 0.041558 | 3725/1843/2817/3326/7132/10000/1003/8878/6383/5598/1514/4790/5970/5296/2353/3554/5290/858/6885/7422/6382/7184/857/5590/9446/ | 35 |

| | | | | | | |
|---|---|---|---|---|---|---|
| | osis | | | | 25828/3162/5327/5602/3551/3791/208/4205/7056/6714 | |
| hsa04012 | ErbB signaling pathway | 23/1495 | 85/7878 | 0.042501 | 3725/9542/1978/817/3265/10000/5595/1027/5063/5296/1399/5290/2002/4609/1026/6777/25/2932/5602/369/208/7039/6714 | 23 |
| hsa04350 | TGF-beta signaling pathway | 25/1495 | 94/7878 | 0.043043 | 1387/100532736/1634/4089/4756/4088/5595/64750/7046/2033/7057/4681/7027/656/26585/6667/7040/4609/4092/5518/7042/3398/60436/1030/3625 | 25 |
| hsa04666 | Fc gamma R-mediated phagocytosis | 25/1495 | 94/7878 | 0.043043 | 3985/56848/10163/4082/10000/8877/10109/5595/50807/5296/1399/5290/10095/65108/3636/9846/5580/5581/8605/208/8976/4651/81873/55616/1785 | 25 |
| hsa04668 | TNF signaling pathway | 29/1495 | 112/7878 | 0.04309 | 3725/602/7132/8837/10000/153090/5595/3726/9586/1435/3659/6376/4790/90993/5970/5296/5606/2353/5290/6885/7186/5602/3551/10488/208/7128/4792/23118/8772 | 29 |
| hsa05161 | Hepatitis B | 40/1495 | 163/7878 | 0.045036 | 1387/3725/3265/10000/4089/4088/5595/7046/9586/2033/4790/148022/90993/5970/5296/5606/2353/5290/2002/7040/6885/4609/7529/1026/6777/581/1017/5602/3551/7042/369/4214/10488/208/4775/353376/4792/23118/6714/8772 | 40 |
| hsa05130 | Pathogenic Escherichia coli infection | 16/1495 | 55/7878 | 0.045456 | 9475/347733/10381/4691/10109/2534/84617/100506658/10095/7280/81027/25/999/8976/10383/81873 | 16 |
| hsa04625 | C-type lectin receptor signaling pathway | 27/1495 | 104/7878 | 0.048193 | 3725/6237/602/8915/3265/10000/5530/5595/5781/23365/3659/4790/3710/5970/5296/5290/22800/4193/5602/5580/3551/208/1540/4775/9261/4792/6714 | 27 |

**Table S1. Enriched DM genes in KEGG pathways in the T2D dataset.** Using the RADAR-detected DM genes in the T2D dataset, we analyzed for enriched KEGG pathways and highlighted a few T2D-related pathways.

### Table S2

| chr | start | end | name | score | strand | thickStart | thickEnd | itemRgb | blockCount | blockSizes | blockStarts | logFC | p_value |
|---|---|---|---|---|---|---|---|---|---|---|---|---|---|
| chr15 | 98648988 | 98649137 | IGF1R | 0 | + | 98649038 | 98649087 | 0 | 1 | 149 | 0 | -0.663 | 2.84e-07 |
| chr4 | 15003223 | 15003372 | CPEB2 | 0 | + | 15003273 | 15003322 | 0 | 1 | 149 | 0 | -1.46 | 1.05e-06 |
| chr17 | 80395488 | 80395637 | RNF213 | 0 | + | 80395538 | 80395587 | 0 | 1 | 149 | 0 | -1.27 | 2.013e-06 |
| chr8 | 143429196 | 143429395 | MAFA | 0 | - | 143429246 | 143429345 | 0 | 1 | 199 | 0 | -0.838 | 5.26e-08 |
| Chr20 | 382532 | 388065 | TRIB3 | 0 | + | 382532 | 388015 | 0 | 2 | 95,55 | 0,5429 | -0.548 | 2.37e-0.5 |

**Table S2. Selected DM sites for experimental validation.** The table shows the peak information of selected putative DM site from RADAR analysis. The genome coordinate is based on hg38. The shown peak table was extended 50bp towards both upstream and downstream to search for RRACH motif match because the RNA molecules for $m_6A$-seq was fragmented to ~150 nt but our sequence reads were only 50bp. Consequently, the estimated peak locations could have position shift from the real peak for up to 100 bp. The extension was intended to take this uncertainty into account.

**Table S3**

| Name | Sequence |
|---|---|
| IGF1R_up | TAG CCA GTA CCG TAG TGC GTG CGC GAC GCA GTT CGC AAG ATC GCC CCG AAG |
| IGF1R_down | /5Phos/CC GGG TCA CAG GCG AGG CCG GCG AGG GGC CAG AGG CTG AGT CGC TGC AT |
| TRIB3_up | TAG CCA GTA CCG TAG TGC GTG AGC AAG ATG CAT AAG TAC CAT CCT TGG GAG |
| TRIB3_down | /5Phos/CT TAG AAA GCT CCC CAG GTT CGA GGC TGG GCA GAG GCT GAG TCG CTG CAT |
| CPEB2_up | TAG CCA GTA CCG TAG TGC GTG AGC GGC GGA GGC GGC GGC GGC GGC TTC GAG |
| CPEB2_down | /5Phos/CC GGA GGG TGG GGA AGG TGG GGA GGG CTG ACA GAG GCT GAG TCG CTG CAT |
| RNF213_up | TAG CCA GTA CCG TAG TGC GTG CCT TCT GAG GCA GAG GTG TAA GCG TTT CAG |
| RNF213_down | /5Phos/CC CAG ATC GGC TAC AGG GAG TGG CGC TCA GCA GAG GCT GAG TCG CTG CAT |
| MAFA_up | TAG CCA GTA CCG TAG TGC GTG GGC CTG GTG TCC ACG TCC TGT ACC GCG GAG |
| MAFA_down | /5Phos/CC GAG CCG AGG CCC CGA GAG GCC TGC GCG ACA GAG GCT GAG TCG CTG CAT |
| PDX1_up | TAG CCA GTA CCG TAG TGC GTG CTA ATT GAA TAC AAG GAG GCA AAT TCT AAG |
| PDX1_down | /5Phos/CT GAA CAG AAT ACA GAA AAT TCT GAC AGT CCA GAG GCT GAG TCG CTG CAT |
| IGF1R_qPCR_F | GCC GCT CAT TCA TTT TGA CT |
| IGF1R_qPCR_R | CTA GGC GAG GAA AAA CAA GC |
| TRIB3_qPCR_F | AAC CTT CAG TGC CTT CCA GA |
| TRIB3_qPCR_R | TGT TGT CAG CTC AAG GAT GC |
| CPEB2_qPCR_F | TTT CCA CCA AAA GGC TAT GC |
| CPEB2_qPCR_R | AGC CCT TAA TGG CCT AGG AA |
| RNF213_qPCR_F | ACA CCT CTG CCT CAG AAG GA |
| RNF213_qPCR_R | TGA AGG GGC ATT TTT AGC AC |
| MAFA_qPCR_F | GCG GAG AAC GGT GAT TTC TA |
| MAFA_qPCR_R | AAG GAA AGG GAG GCT GAG AA |
| PDX1_qPCR_F | AGC AGT GCA AGA GTC CCT GT |
| PDX1_qPCR_R | CAC AGC CTC TAC CTC GGA AC |

**Table S3. Oligo probes sequences and qPCR primer sequences.** We designed an up and a down probe flanking the putative DM $m_6A$ site leaving the $m_6A$ nucleotide as a gap. For each pair of oligo probes, we designed an overhanging universal primer sequence at their 5' and 3' end, respectively. The table shows the sequence of oligo probes we used as well as the qPCR primers we used to quantify gene level variation.
